# Directionality of PYD filament growth determined by the transition of NLRP3 nucleation seeds to ASC elongation

**DOI:** 10.1101/2021.11.25.470035

**Authors:** Inga V. Hochheiser, Heide Behrmann, Gregor Hagelueken, Juan F. Rodríguez-Alcázar, Anja Kopp, Eicke Latz, Elmar Behrmann, Matthias Geyer

## Abstract

Inflammasomes sense intrinsic and extrinsic danger signals to trigger inflammatory responses and pyroptotic cell death. Homotypic pyrin domain (PYD) interactions of inflammasome forming Nod-like receptors with the adaptor protein ASC mediate oligomerization into helical filamentous assemblies. These supramolecular organizing centers recruit and activate caspase-1, which results in IL-1β family cytokine maturation and pyroptotic cell death. The molecular details of the critical step in signal transduction of inflammasome signaling, however, remain ill-defined. Here, we describe the cryo-EM structure of the human NLRP3 PYD filament at 3.6 Å resolution. We identify a unique pattern of highly polar interface residues that form the homomeric interactions leading to characteristic filament ends that we designate as A- and B-end, respectively. Coupling a titration polymerization assay to cryo-EM, we demonstrate that the ASC adaptor protein elongation on NLRP3 PYD filament seeds is unidirectional, associating exclusively to the B-end of the NLRP3 filament. Notably, NLRP3 and ASC PYD filaments exhibit the same symmetry in rotation and axial rise per subunit, allowing for a continuous transition between NLRP3 as the nucleation seed and ASC as the elongator. Integrating the directionality of filament growth, we present a molecular model of the ASC speck consisting of active NLRP3–NEK7, ASC, and Caspase-1 proteins.

## INTRODUCTION

Innate immune cells are able to recognize invading pathogens or cellular damage by germline-encoded pattern recognition receptors (PRRs) (Takeuchi and Akira, 2010). Among these, the AIM2-like receptors (ALRs) and NOD-like receptors (NLRs) form supramolecular complexes termed inflammasomes, which regulate the activation of highly pro-inflammatory cytokines and pyroptotic cell death (Martinon et al., 2002). These large cytosolic assemblies consist of sensor, adaptor, and effector proteins and form within minutes after recognizing their specific triggers (Broz and Dixit, 2016; Swanson et al., 2019).

Canonical inflammasome formation starts with the activation-induced oligomerization of ALR or NLR sensor proteins that, in turn, recruit pro-caspase-1, either by direct homotypic protein domain interactions or via the adaptor protein ASC (apoptosis-associated speck-like protein containing a CARD). Proximity-induced autoproteolytic cleavage yields active caspase-1 that mediates activation and release of highly pro-inflammatory cytokines of the IL-1 family and triggers a pro-inflammatory form of cell death called pyroptosis (Franchi et al., 2012; Latz et al., 2013). NLRP3 is the best-studied inflammasome protein to date as it is involved in the recognition of a plethora of different activating ligands encountered during bacterial, viral or fungal infections, as well as in sterile inflammation (Swanson et al., 2019). While during pathogen encounter, NLRP3 activation may be a beneficial event of the innate immune response, it confers detrimental effects during sterile inflammation, manifesting in autoinflammatory diseases like atherosclerosis, gout, and Alzheimer’s disease (Friker et al., 2020; Ising et al., 2019; Strowig et al., 2012; Venegas et al., 2017). Moreover, several mutations in the *nlrp3* gene that induce the undue activation of the NLRP3 inflammasome were linked to a group of rare, inherited, autoinflammatory diseases, which are summarized as cryopyrin-associated periodic syndromes (CAPS) (Hoffman et al., 2001).

NLRP3 inflammasome formation is thought to be a two-step process. In the first priming event, NLRP3 expression is upregulated via nuclear factor-kB (NF-kB) dependent signaling (Bauernfeind et al., 2009). At this stage, NLRP3 is present in an autoinhibited but signaling competent conformational state. The engagement of an activating signal then initiates the second step of NLRP3 inflammasome formation (Swanson et al., 2019). Activation-driven conformational changes allow for the oligomerization of NLRP3 with the participation of its central NACHT domain and the N-terminal Pyrin domain (PYD). Oligomerized NLRP3 recruits the bipartite adaptor protein ASC through homotypic PYD–PYD domain interactions, serving as a nucleation platform for the formation of filamentous ASC assemblies (Lu et al., 2014; Sborgi et al., 2015).These prion-like structures then cluster into large ASC specks that trigger caspase-1 activation (Baroja-Mazo et al., 2014; Cai et al., 2014; Franklin et al., 2014). CARD–CARD domain interactions mediate this assembly that bundles multiple ASC filaments into a large conglomerate called ASC speck and also perform heteromeric interactions between the ASC and the Caspase-1 CARDs (Dick et al., 2016). This self-propagating process based on a cascade of ordered interactions allows for a robust signal amplification mechanism (Dick et al., 2016; Ruland, 2014).

To date, two ASC-dependent inflammasome sensors, namely AIM2 and NLRP6, have been shown to form PYD filaments that can nucleate ASC polymerization *in vitro* and in cells (Lu et al., 2015; Shen et al., 2019). For NLRP3, previous *in vitro* studies suggested the need for the NACHT domain for successful ASC polymerization (Lu et al., 2014), while the PYD alone was sufficient to induce ASC speck formation in HEK293T cells (Marleaux et al., 2020; Stutz et al., 2017). A filament structure of the PYD of the adaptor protein ASC has been determined previously (Lu et al., 2014; Sborgi et al., 2015), and a model for ASC filament formation with the CARD domain serving as a signal amplification hub for inflammasome assembly has been proposed (Dick et al., 2016). However, the critical step necessary for inflammasome signal transduction, the PYD filament interaction between the inflammasome sensor and the adaptor ASC, remains poorly understood.

In this study, we determined the cryo-EM structure of the human NLRP3^PYD^ filament at 3.6 Å resolution, identifying the same symmetry in rotation and axial rise per subunit compared to the ASC^PYD^ filament (Lu et al., 2014). Each NLRP3^PYD^ subunit is arranged in a hexagon-like assembly creating three asymmetric interfaces that interact with six adjacent PYDs. The complementary interfaces form unique surfaces at the filament ends, designated as A- and B-end, respectively. Importantly, we developed an *in vitro* filament reconstitution assay that allows for the homotypic transition from an NLRP3 PYD nucleation seed to ASC filament elongation. Using cryo-EM, we find that filament transition is unidirectional and exclusively occurs at the B-end of the NLRP3^PYD^ filament. This defines the filament growth direction of ASC adaptor elongation, the assembly of ASC^CARD^ domains into filament bundles for ASC specking, and the interactions with pro-caspase-1 for its activation by auto-proteolysis. Our observations reveal the dynamics of homotypic filament formation and have implications for the possible interference by antibodies or small molecules at the filament growing site.

## RESULTS

### Polymerization of recombinant NLRP3^PYD^ and cryo-EM structure determination

For structural analyses, human NLRP3^PYD^ was purified with a TEV cleavable N-terminal GST-tag from *E. coli* cells. The PYD filament formation was induced by removing the affinity tag, followed by incubation of the gel-filtered NLRP3^PYD^ protein at 37°C, which resulted in long and straight individual filaments suitable for cryo-EM structure determination (Figures 1A, S1A and S1B). Micrographs were recorded with a Titan Krios transmission electron microscope, equipped with a Falcon 2 camera. 3D refinement, using the helical parameters of the ASC^PYD^ filament (Lu et al., 2014) (PDB 3J63) as a starting point, yielded an electron density map with a resolution of 3.6 Å (Figures S1C-F). Rigid body fitting of the NLRP3^PYD^ crystal structure (Bae and Park, 2011) (3QF2) into the cryo-EM map was followed by real-space refinement to achieve the final NLRP3^PYD^ filament structure (Figure 1B). The final model encompasses residues 3-94, exhibiting an RMSD-value of 0.54 Å with the crystal structure of NLRP3^PYD^ (Figure 1C). Within the six helical bundle of the PYD (helices α1 to α6), only slight deviations between the filament and the globular structure are seen in the region at the end of helix α3 and the beginning of helix α4, whereas the overall arrangement is well preserved (Figure S2).

**Figure 1.**
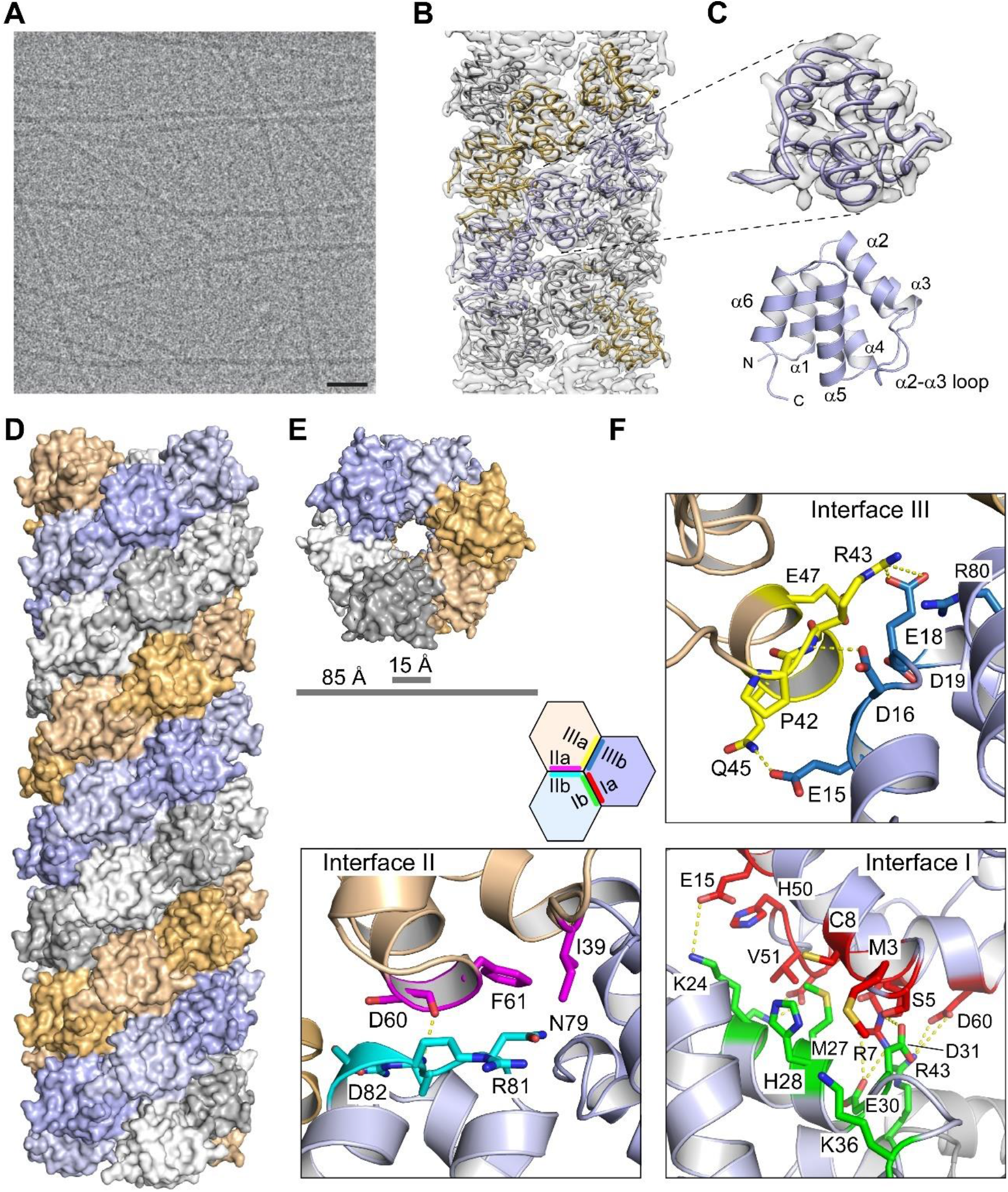
Cryo-EM structure of the NLRP3^PYD^ filament. (A) Representative cryo-EM micrograph of NLRP3^PYD^ sample after incubation at 37°C overnight. The scale bar corresponds to 50 nm. (B) Superposition of the NLRP3^PYD^ filament model with the reconstructed electron density. (C) Close up of a single NLRP3^PYD^ filament chain in the electron density map and ribbon representation of the NLRP3^PYD^ structure. (D) Surface representation of the side view of the NLRP3^PYD^ filament with three-start helical symmetry. (E) Top view of the NLRP3^PYD^ filament showing the three-start helical symmetry. (F) Detailed interactions in the type I, type II and type III interfaces of the NLRP3^PYD^ filament. A schematic diagram of the hexagonal assembly of the NLRP3^PYD^ domain within the filament structure indicates the three asymmetric interfaces.

### Architecture and interfaces of the NLRP3^PYD^ filament

The NLRP3^PYD^ filament is a cylindrical, hollow structure with an outer and inner diameter of ∼85 Å and ∼15 Å, respectively (Figures 1D and 1E). It displays a three-start helical symmetry of 54.9° right-handed rotation and an axial rise of 14.3 Å per subunit. The filament assembles as a triplicate by the C*_n_*(3) helical symmetry. As for other death domain folds (Ferrao and Wu, 2012), the NLRP3^PYD^ filament is composed of three major asymmetric interfaces, designated as type I, II, and III interfaces, with the opposing interacting surfaces named as a and b, respectively. In the NLRP3^PYD^ filament interface I is the largest in size, burying about 1020 Å^2^ of surface area (a and b sides together) on one PYD molecule. This interface, which is also the most dominant, is composed of polar residues that appear mostly conserved within the PYD family (Park, 2012). The type II and III interfaces are more variable and bury about 430 and 500 Å^2^ surface areas in the NLRP3^PYD^ filament. Due to the three-start helical symmetry, the type I interface mediates intra-strand interactions, whereas the type II and type III interfaces mediate inter-strand interactions (Figure 1D). All three interfaces are constituted through electrostatic, polar, as well as hydrophobic interactions.

Interface I is mediated by interactions between residues of the α1 and α4 helix of one chain with residues of the α2 and α3 helices of the neighboring chain (Figure 1F). Within a distance of 4 Å, there is a remarkable high number of eight salt bridges, involving each four charged residues on both interface sites, as well as five hydrogen bonds constituting this interface. Here, the guanidinium group of Arg7 is most prominent, forming tight electrostatic interactions with the side chains of Glu30 and Asp31. Interface II is mediated by interactions of α4 residues of one chain with the α5-α6 loop of a neighboring strand chain (Figure 1F). Yet, only one hydrogen bond at 3.1 Å distance is formed between the backbone carbonyl group of Asp60 and the backbone amine of Arg81. Finally, interactions between helix α3 of one chain with the loop of helix α1-α2 of the neighboring strand chain form the type III interface (Figure 1F). Two salt bridges between Arg43 and Glu18 and three hydrogen bonds constitute the binding besides many hydrophobic interactions. Unexpectedly, with 430 Å^2^ the buried surface area of interface II appears smaller than the concealed surface area of interface III with 500 Å^2^ in the NLRP3^PYD^ filament structure. This is in marked difference to the ASC^PYD^ filament, where interface II and III each bury 540 and 360 Å^2^ of surface area, respectively (Lu et al., 2014). Besides, interactions between three subunits within the filament at the interface tips appear characteristic for the NLRP3^PYD^ assembly as multiple residues undergo interactions with more than one interface.

### Tripartite interactions at the interface tips

The hexagonal assembly in the filament structure where every PYD molecule is surrounded by six other PYD molecules contains two unique corner sites. There is one remarkable charged interface at the joints of three molecules that appears unique to the NLRP3^PYD^ filament. This interface involves three arginines in a staggered arrangement from three different interface sites, with Arg43_IIIa_ from one molecule occupying the central position, headed by Arg7_Ia_ from a second molecule at one site and Arg80_IIb_ from a third molecule at the other site (Figure 2A). The assembly of these three positively charged residues is facilitated by the compensation with six negatively charged residues in close vicinity to the arginines. A mixed salt bridge network from three different molecules is mediated by Asp60_IIa_ directly facing Arg43_IIIa_ and backed up by Glu18_IIIb_ (Figure 2B). Glu30_Ib_ and Asp31_Ib_ from one molecule are close to Arg7_Ia_ from a second molecule, and Arg80_IIb_ is finally interacting with Asp16_IIIb_ and Asp82_IIb_ within one molecule. The tripartite interaction in the filament assembly between Asp60, Arg43 and Glu18, however, from three different PYD molecules is unique to NLRP3 within the known structures of filament-forming PYDs of the inflammasome as no other PYD sequence contains this charge network (Figure 2A).

**Figure 2.**
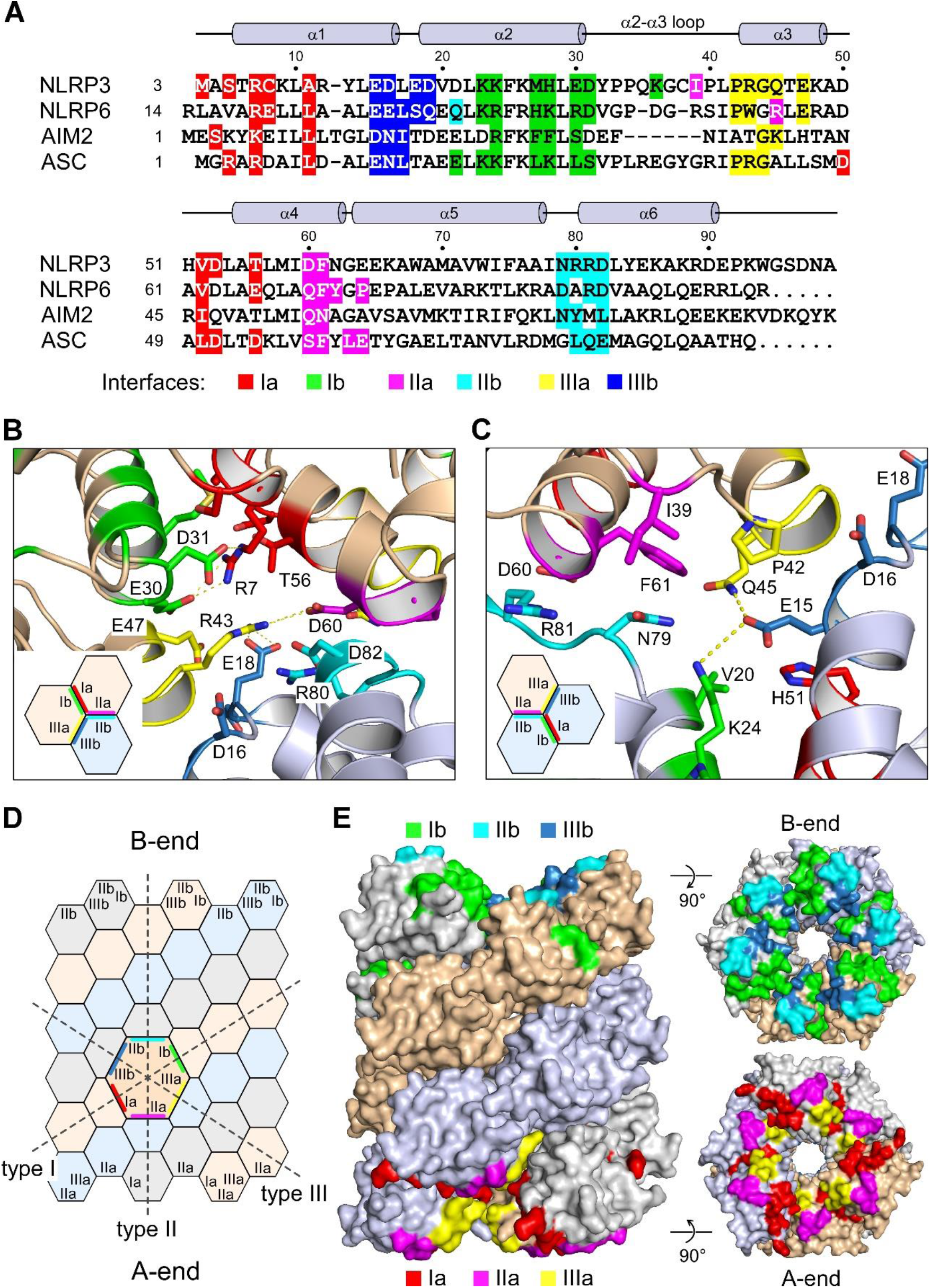
Sequence conservation and interface sides in the NLRP3^PYD^ filament. (A) Structure-based sequence alignment of the four PYDs whose filament structures have been determined. Interface forming residues are highlighted and shown below the secondary structure elements of NLRP3^PYD^. Residues are colored according to the respective interface sites as determined for the PYD filaments of NLRP3 (this study; 3-99), NLRP6 (6NCV, 14-104), AIM2 (6MB2, 1-94), and ASC (3J63, 1-91). (B) Tripartite interactions at the tips of the interfaces. Positively and negatively charged residues of all six interface sites assemble to a highly polar interaction. (C) Close up of the second unique interaction at the interface tips. (D) Schematic diagram of the hexagonal subunit assembly in the three-start helix and designation of the interface sites. The a- and b-sides of the three asymmetric interfaces assemble at the filament’s ends to characteristic surfaces, designated as A- and B-end of the filament, respectively. (E) Surface representation of the NLRP3^PYD^ filament model with the interfaces at the top and bottom layer colored according to the scheme in (A). The a- and b-interfaces merge at the filament ends to the A- and B-end of the filament.

The second unique corner in the hexagon assembly of the NLRP3^PYD^ filament structure is less pronounced but involves similarly the tripartite formation of a hydrogen bond network. Here, residues from five interface sites come together with Glu15_IIIb_ in an exposed position close to Lys24_Ib_ from a second molecule and Gln45_IIIa_ from a third molecule (Figure 2C). Asn79_IIb_ from the second molecule and hydrophobic interactions with Phe61_IIa_ and Pro42_IIIa_ from the third molecule complement these homotypic interactions at the tip of the three protein interfaces. The three types of asymmetric interfaces with the opposing interactions sites a and b in the honeybee comb-like assembly of PYD subunits assemble into one continuous surface at the two ends of the filament (Figure 2D). We therefore define these sites as the A-end and B-end of the NLRP3^PYD^ filament, respectively (Figure 2E).

### Mutational analysis of interface forming residues

To study the impact of individual residues on the filament formation of NLRP3^PYD^, we performed site-directed mutagenesis analyses with recombinant proteins and in cells. Dynamic light scattering (DLS) experiments and negative-stain EM images were recorded to visualize filament formation. Based on the filament structure, seven residues from five different interface sites were selected, and the charge reversal mutations R7E, E15R, R43E, R80E, R81E, the double mutation K23E/K24E, and the mutation M27E introduced. In addition, the four NLRP3^PYD^ CAPS mutants D19H, D31V, H51R, and A77V were analyzed that are known for their phenotype in Muckle-Wells disease (Hu et al., 2017; Martorana et al., 2017; Turunen et al., 2018). First, the kinetic profile of NLRP3^PYD^ filament formation was monitored in a time-course experiment using DLS. At a protein concentration of 48 µM and a temperature of 25°C, the saturation of the oligomerization process was achieved in about 40 min as revealed by the normalized growth signal (Figure 3A). The polymerization status of the NLRP3^PYD^ interface and CAPS mutants was determined at a time point of 90 min when the polymerization had reached a stable plateau (Figure 3B). All of the tested interface mutants lost their ability to oligomerize into ordered polymers but instead formed unspecific aggregates (Figure 3C), confirming the importance of each mutated residue for the homotypic filament formation. In contrast, of the tested CAPS mutants, only the D31V variant lost the ability to polymerize, while the A77V mutant formed short but ordered filaments (Figures 3B and 3C). The disease mutations D19H and H51R showed a strong polymerization behavior, which made an analysis by gel filtration and subsequent DLS impossible. However, this phenotype was confirmed by negative-stain EM analysis.

**Figure 3.**
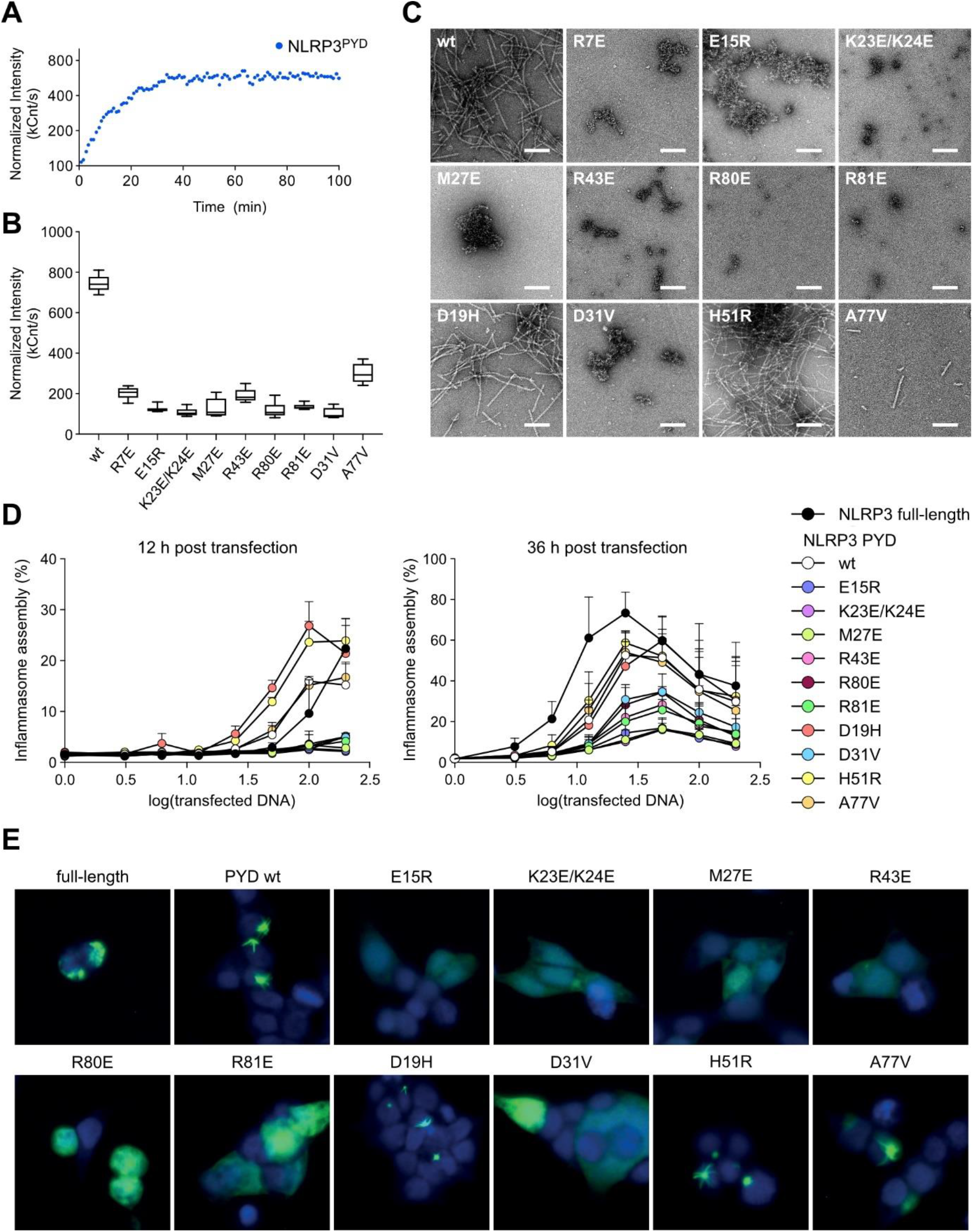
Mutational analysis of NLRP3^PYD^ interface residues. (A) NLRP3^PYD^ polymerization *in vitro* monitored by dynamic light scattering. Normalized intensity signals are recorded as a function of time and shown here for one representative experiment. (B) Polymerization status of NLRP3^PYD^ and mutants carrying additional interface and CAPS mutations monitored by dynamic light scattering after an incubation period of 1.5 h at 37°C. Normalized intensity signals are reported for each variant (n=3). (C) Negative-stain electron micrographs of recombinant NLRP3^PYD^ and interface disrupting mutants (upper two lines) or CAPS mutants (lower line). CAPS mutations D19H and H51R even increase PYD filament formation. (D) Quantification of HeLa cells stably expressing ASC-mTurquoise transfected with full-length NLRP3-mCitrine, NLRP3^PYD^-mCitrine or indicated NLRP3^PYD^ mutants, 12 h and 32 h after transfection. Dots are representative of two (n=2±SD; 12 h) and five (n=5±SEM; 32 h) independent experiments for the two different time points, respectively. (E) Images of HeLa cells stably expressing ASC-mTurquoise transfected with NLRP3-mCitrine fusion constructs as used in (D).

To test the implications of the interface mutants in a physiological context, we performed ASC speck assays in HeLa cells. Here, full length, wild type NLRP3-mCitrine or wild type or mutant NLRP3^PYD^ with mCitrine fused C-terminally to the protein were transfected into HeLa cells stably expressing ASC-mTurquoise. Priming of ASC polymerization by a filament nucleation factor induces the ASC speck formation that can be subsequently quantified (Stutz et al., 2017). As a readout, the percentage of cells that develop ASC specks was determined in dependence on the amount of transfected DNA and analyzed for two different time points (Figures 3D and 3E). The NLRP3^PYD^ protein stably triggered ASC speck formation, albeit to a lesser extent than full-length NLRP3. The mutants E15R, K23E/K24E and M27E most successfully abolished the ability of the NLRP3^PYD^ to nucleate ASC polymerization. The mutants D31V, R43E, R80E and R81E also impaired ASC speck formation, however, to a lesser extent, suggesting only partial defectiveness. ASC specking ability of the CAPS mutant A77V was similar to the wild type protein, whereas the CAPS mutants D19H and H51R showed an even increased efficacy of nucleating ASC. Fluorescence images showing the speck formation of full-length NLRP3 and NLRP3^PYD^ wild type and mutant protein are shown in Figures 3E and S3.

### The transition of the NLRP3 to ASC PYD filament is unidirectional

With the structure of the NLRP3^PYD^ filament determined, we next aimed at analyzing the transition to ASC adaptor protein elongation on a molecular level. ASC polymerization is triggered by NLRP3 nucleation seeds (Dick et al., 2016; Lu et al., 2014), leading ultimately to the formation of large ASC specks. These specks are thought to arise from a continuous transition of NLRP3 and ASC PYD filaments and subsequent crosslinking of ASC^PYD^ stems via CARD–CARD domain interactions (Dick et al., 2016; Schmidt et al., 2016). Seeding of ASC filaments by NLRP3 *in vitro* was achieved by developing a polymerization protocol, where 50 µl of 50 µM monomeric NLRP3^PYD^ at pH 3.8 were rapidly neutralized by the addition of 1 µl 3 M Tris buffer (pH 8) to initialize filament growth (Figure 4A). After 3 min incubation time at 25°C, monomeric, soluble ASC-mCherry at 16 µM concentration (pH 3.8) was added at a 1:50 molar ratio of ASC to NLRP3 and incubated for 5 min. This step was repeated twice to slowly raise the concentration of ASC and allow elongation of the NLRP3^PYD^ seeded ASC-mCherry filaments. To distinguish the two filament types, we used a 50 kDa ASC-mCherry fusion protein that is fourfold larger than the 12 kDa NLRP3^PYD^ protein and thus appeared markedly thicker on the negative stain micrographs compared to the thinner NLRP3^PYD^ filaments (Figure 4B). Indeed, transitions of the NLRP3^PYD^ nucleation seeds to ASC filament elongation were observed in many instances (Figure 4C). Following the gentle titration procedure that leads to a very dilute ASC concentration initially, only a few cases of spontaneous ASC polymerization events were seen, whereas in approximately half of the cases elongation of ASC on NLRP3^PYD^ filaments occurred. Intriguingly, ASC filament growth was observed only on one end of the NLRP3^PYD^ filament but never on both ends, suggesting a directionality in the transition from the NLRP3 to ASC filament. Repeated titration of ASC-mCherry to the solution led to longer ASC filament growth, indicating a preference for the elongation of existing filaments compared to spontaneous *de novo* polymerization of ASC alone (Figure 4D). Of note, the thicker ASC-mCherry filaments are always centered on the NLRL3^PYD^ seeds, indicating that the transition is mediated by the N-terminal PYD of the ASC protein. The congruent helical symmetry between filaments assembled by AIM2^PYD^ and the downstream adaptor ASC^PYD^ has been already elegantly shown by the Egelman and Sohn laboratories (Matyszewski et al., 2021; Morrone et al., 2015).

**Figure 4.**
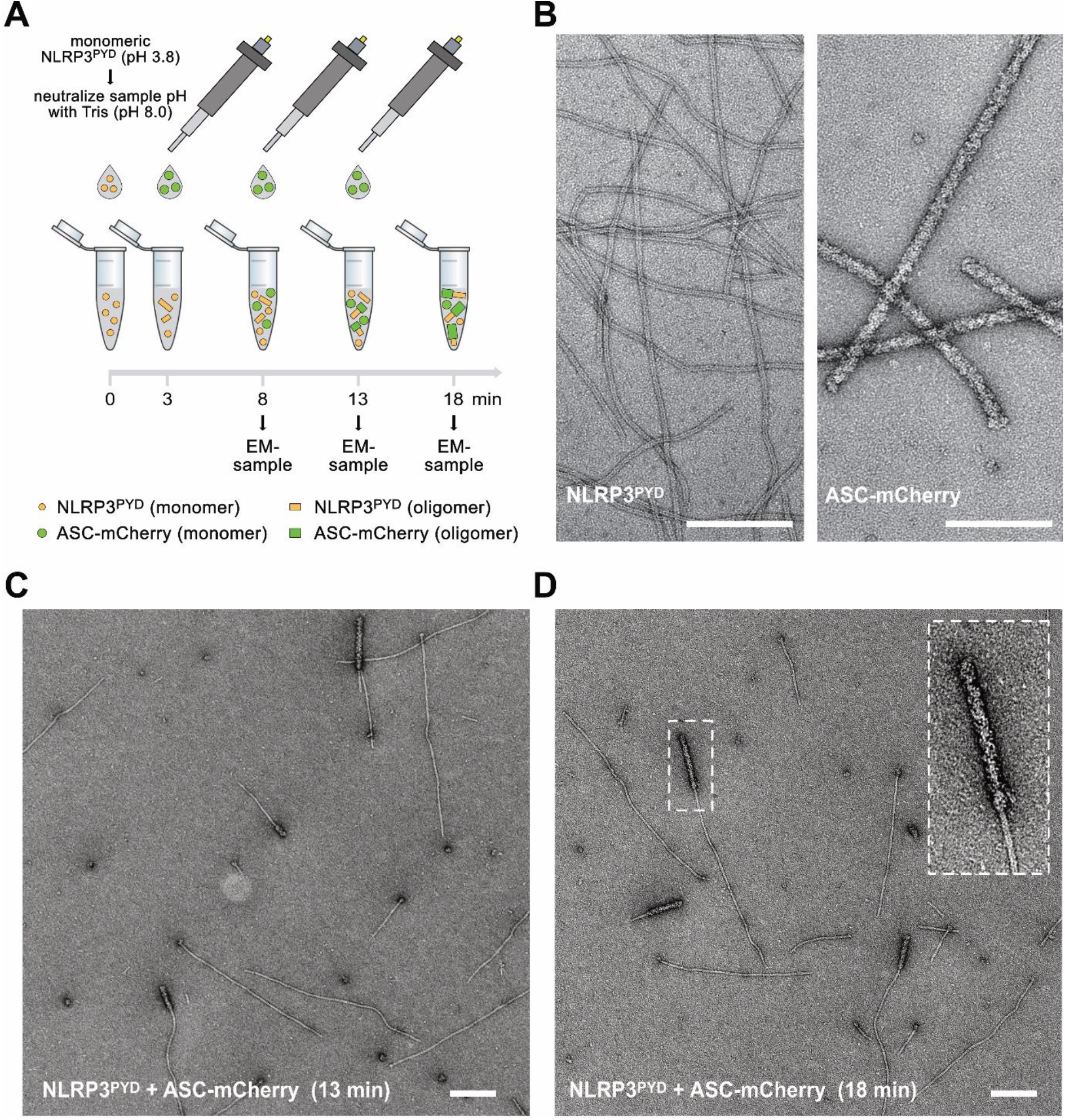
ASC filament polymerization on NLRP3^PYD^ nucleation seeds. (A) Cartoon depiction of an NLRP3^PYD^ induced ASC filament polymerization assay. Filament formation of monomeric NLRP3^PYD^ is induced by neutralization to pH 8.0. Stepwise titration of highly diluted ASC-mCherry leads to the elongation on NLRP3 filament seeds. (B) Negative-stain electron micrographs of NLRP3^PYD^ (left) and ASC-mCherry filaments (right). The scale bar corresponds to 200 nm. (C) Elongation of ASC-mCherry filaments on NLRP3^PYD^ filament seeds after the first two titration steps. Several thin NLRP3^PYD^ filaments are seen out of which thicker ASC-mCherry filaments grow out. Note that at the most, one transition is seen from an NLRP3 to an ASC filament, suggesting a directionality of filament growth. (D) Elongation of the ASC-mCherry filament on NLRP3^PYD^ seeds after three titration steps. Increasing the ASC concentration induces ASC filaments to grow longer on the NLRP3^PYD^ nucleation seeds.

### ASC elongates at the B-end of the NLRP3 PYD filament

To identify the site of elongation on the NLRP3^PYD^ filament and the directionality of ASC filament growth, we performed cryo-EM imaging, providing the resolution required for the determination of filament directionality. The titration protocol for the domain transition was adjusted to generate longer NLRP3^PYD^ filaments that allowed for the analysis of straight filament stretches. In the cryo-EM micrographs, ASC-mCherry filaments appear more diffuse than in the negative-stain mode; nonetheless, they could be clearly identified (Figure 5A). Only NLRP3^PYD^ filaments merging into ASC-mCherry filament transitions were selected and used for subsequent image analysis. Average class sums calculated from 19,351 particles that were selected from 423 micrographs unambiguously revealed a saw-tooth shaped density, reflecting the directionality of the filaments. A selection of average class sums showing this characteristic saw-tooth pattern is displayed in Figure 5B. The electron density map of the filament confirms this silhouette, indicated by the black lines lining the envelope (Figure 5C). Superposition of our filament structure onto those classes reveals the orientation of the filament in the average sums as the saw-tooth shaped density arises from the bending of the C-terminal helix α6 of the PYD (Figure 5D). It should be noted that this characteristic pattern must not necessarily be visible in the average class sums as the pitch of the PYD in the three-start helix leads to various projections within the short sections used for object averaging.

**Figure 5.**
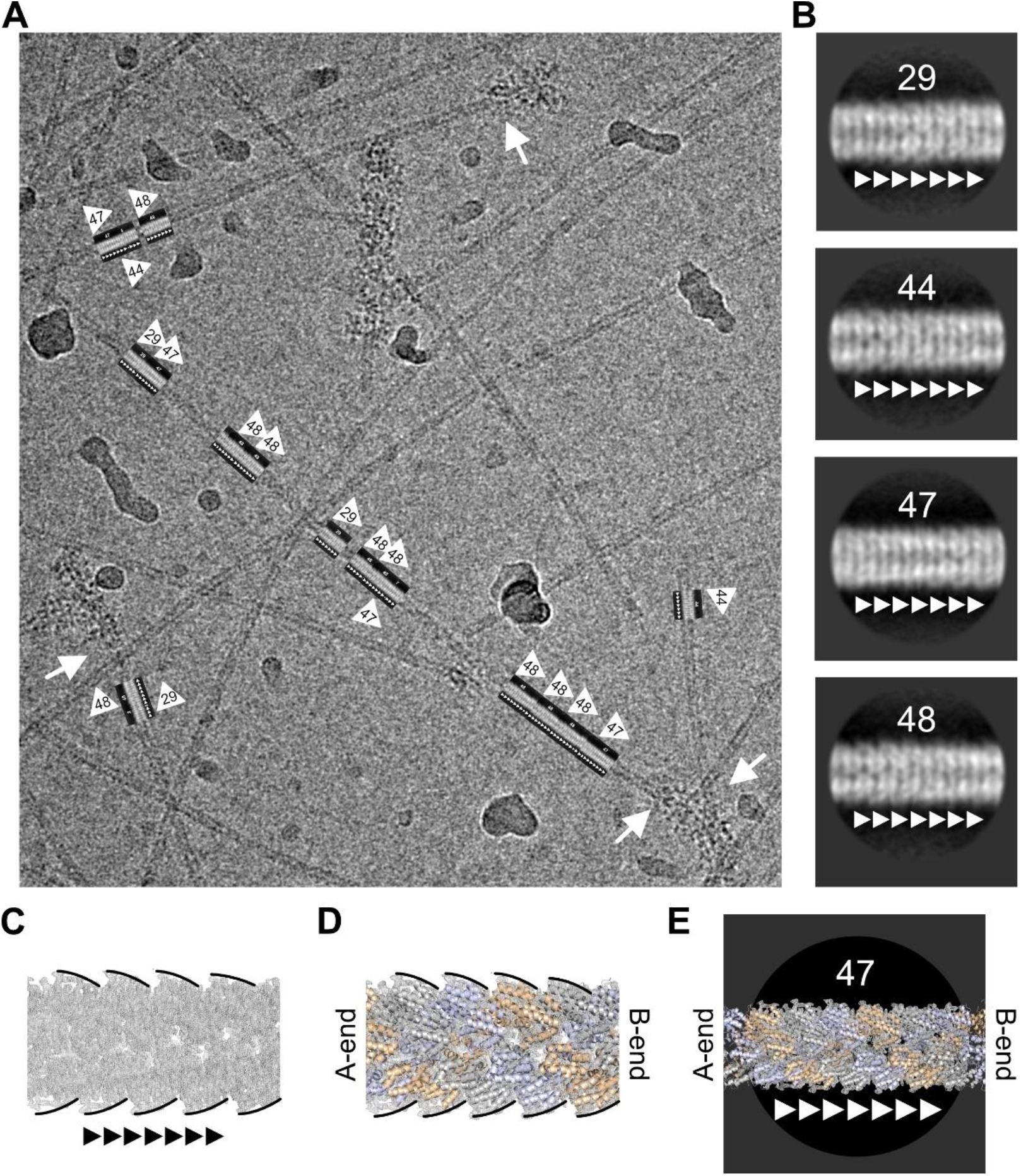
ASC exclusively elongates at the B-end of the NLRP3^PYD^ filament. (A) Representative cryo-EM micrograph of NLRP3^PYD^ filament to ASC-mCherry filament transitions. Long NLRP3^PYD^ filaments were grown that allow for analysis of straight segments. The direction of the transition site to ASC-mCherry is indicated by small arrows in the boxes. Transition sites of NLRP3 to ASC filaments are indicated by white arrows. (B) Average class sums of individual segment particles for four representative classes. The direction towards the NLRP3 to ASC PYD transition is indicated by small arrows. A saw tooth pattern at the edges and a perforation pattern in the filament center is characteristic for some classes. (C) Electron density map of the NLRP3^PYD^ filament with the characteristic saw tooth pattern identifies the A- and B-end of the filament. (D) Modelling of the filament ribbon structure into the electron density map. (E) Overlay of the filament structure of the average class sums identifies the directionality of the NLRP3 to ASC PYD filament transition. Shown for the average class 47 the characteristic saw tooth pattern and the perforation identifies the B-end of the NLRL3 filament for the elongation with ASC.

For classes 29, 44, 47 and 48 a superposition allowed to unambiguously correlate the NLRP3^PYD^ filament direction with the given orientation of the class sums. This is shown as an example for class 47, where the left and the right end could be assigned to the A- and B-end of the filament, respectively (Figure 5E). To pin down the directionality of the NLRP3^PYD^ filament seen in the micrographs, we first added a directionality flag into the class averages (white triangles). We then extracted the rotation and translation information for each of the underlying particles (Figures S4 and S5). We used this information to superimpose the flagged class averages onto the original micrographs. This procedure revealed that the white triangles on the picked particles re-assigned to the original micrographs always pointed towards the transition to the ASC filament (Figure 5A). Out of eight average class sums combined in a total of 560 boxed filament sections, none pointed in the opposing direction (Figure S4D). From these data analyses, we conclude that the transition from NLRP3^PYD^ filament seeds to ASC filament elongation exclusively occurs at the B-end of the existing NLRP3^PYD^ filament and that the direction of ASC filament growth is from the A- to the B-end.

### Interactions in the NLRP3–ASC PYD transition interface

Knowing the ASC elongation site on NLRP3 nucleation seeds, we set out to investigate the transition interface between these two homotypic PYD filaments. Twelve subunits of NLRP3^PYD^ and the ASC^PYD^ filament (Lu et al., 2014) were superimposed and aligned according to the growth direction (Figure 6A). As the filaments exhibit the same rotational symmetry and axial rise, a continuous transition is facilitated. Using the GROMACS molecular dynamics package (Van Der Spoel et al., 2005), we optimized the binding interface between the B-end of the NLRP3^PYD^ filament and the A-end of the ASC^PYD^ filament elongation by energy minimization of a fully hydrated protein complex. The interfaces fit very well with complementary charge and hydrogen bond interactions and no atomic clashes (Figures 6B–D). In total, 33 residues constitute the hetero-PYD filament interface, out of which 12 are identical and another nine are similar in both ASC and NLRP3 domains (Figure 6C). Overall, the sequence homology between the ASC^PYD^ and the NLRP3^PYD^ is unexpectedly low with a sequence identity of 22% and a similarity of 43%. Interestingly, residues in interface I are particularly well conserved whereas interface II residues on the b-side diverge markedly between both PYDs.

**Figure 6.**
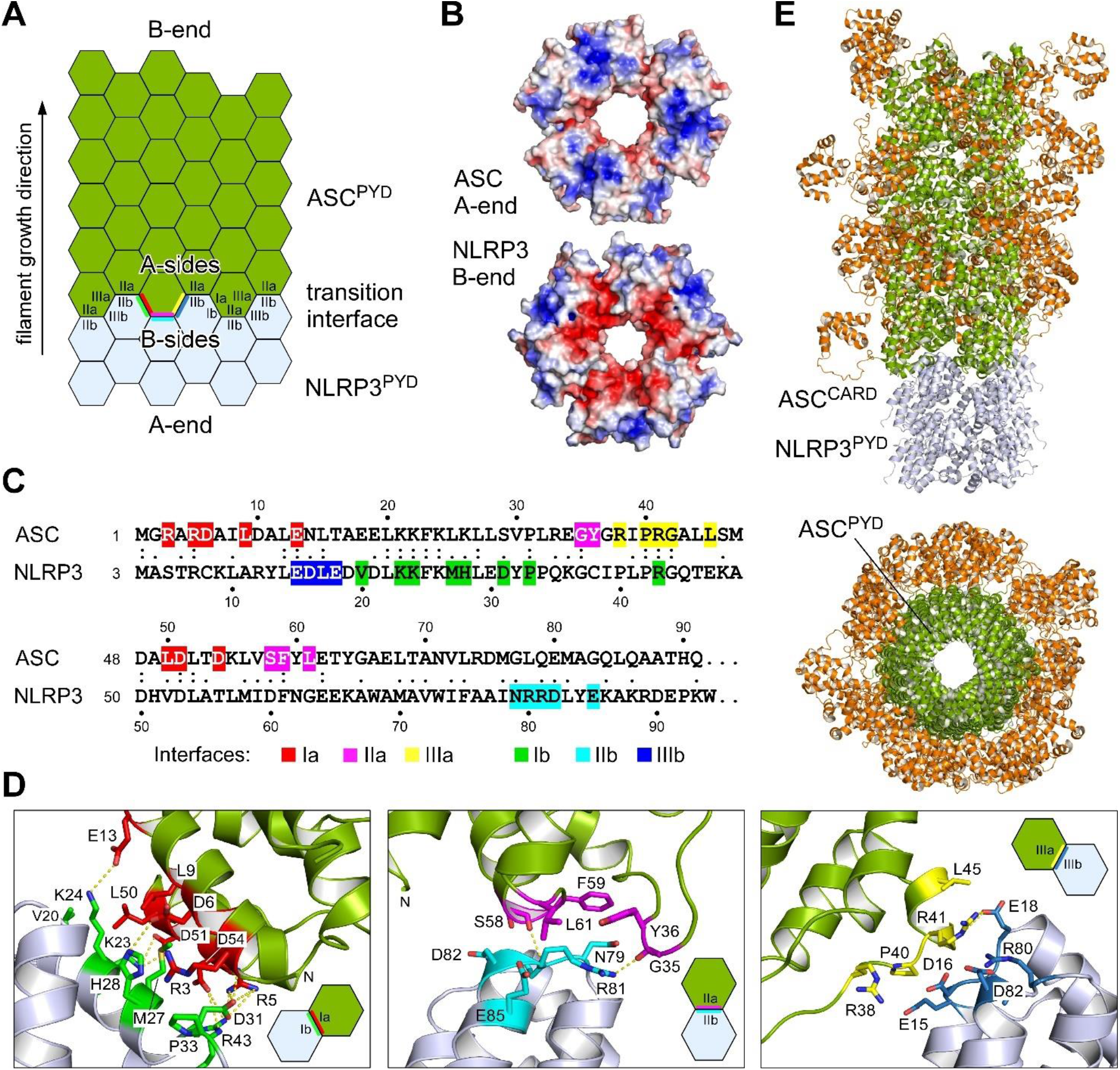
Transition of NLRP3^PYD^ nucleation seeds to ASC^PYD^ filament elongation. (A) Cartoon of the filament transition from the nucleation seed NLRP3^PYD^ to the elongation adaptor ASC^PYD^. The transition interface of the heteromeric PYD interactions are indicated. (B) Electrostatic surface potential of the complementary filaments from the ASC^PYD^ A-end and the NLRP3^PYD^ B-end, displayed from −4 *k*_B_T (red) to +4 *k*_B_T (blue). (C) Sequence alignment of the ASC and NLRP3 PYDs with the interface residues highlighted. Interacting residues from the a side interfaces in ASC and the b side interfaces in NLRP3 are colored as before. (D) Close-up of the interactions in the three interface types I, II and III for the energy minimized transition interface between the NLRP3^PYD^ filament (light blue) and the ASC^PYD^ filament (green). Salt bridges (type I and III) and hydrogen bonds (type II) are indicated by dashed lines. (E) Model of an ASC filament (green) elongating on the NLRP3^PYD^ filament nucleation seed (light blue). Whereas the PYD domain of ASC forms the filamentous stem structure, the CARD domain could adopt variable compositions at the surface of the filament.

The interactions within the heteromeric PYD assembly are visualized for all three interface types in Figure 6D. The entire buried surface area in this transition segment comprises 930 Å^2^ for interface I followed by 560 and 430 Å^2^ for interfaces II and III, respectively. The increase in interface II compared to the NLRP3^PYD^ filament alone (430 Å^2^) results from the a-side residue L61 in ASC, which is a small glycine in NLRP3, and a novel interaction of R81 in NLRP3 with residues G35 and Y36 of the α2-α3 loop in ASC forming a hydrogen-bond with the G35 carboxyl group (Figure 6D). Interestingly, this interaction has no correspondent in NLRP3 due to a different conformation and sequence composition of this loop. A similar interaction is also not seen in the ASC^PYD^ filament, as the corresponding residue Q79 does not reach out as far as R81 in NLRP3. The increase in interface area III compared to the ASC^PYD^ filament alone (320 Å^2^) instead results from b-side residues E18 and R80 in NLRP3, which correspond to small residues T16 and L78 in ASC. From the interface analysis we did not identify a unique restriction mechanism that would prohibit elongation at the filament A-end of NLRP3 with the B-end of ASC. Instead, we speculate that the kinetics of filament growth drives the direction from the A-end to the B-end and thus determines the elongation directionality. To create a model of the NLRP3^PYD^ nucleation seed to ASC adaptor transition, the NLRP3^PYD^ filament was elongated with 32 ASC^PYD^ subunits which each were overlayed with a randomly selected NMR model of the full-length ASC protein (2KN6; de Alba, 2009). This model provides a first glimpse on how the adaptor ASC could bridge from the NLRP3 inflammation sensor to the Caspase-1 pyroptosis effector (Figure 6E).

## DISCUSSION

The formation of an ASC speck is the hallmark of inflammasome activation in immune cells and the signal that initiates pyroptotic cell death. A cascade of sensor, adaptor, and effector protein interactions transmits and amplifies the signal of danger- or pathogen-induced triggers that ultimately leads to cytokine release as a systemic inflammatory response (Broz and Dixit, 2016; Martinon et al., 2002). The human NLPR3 protein is one critical sensor of the innate immune system that induces ASC speck formation upon activation. Here, we show that the Pyrin domain of NLRP3 can assemble into helical filaments similarly as it has been described for the AIM2, NLRP6 and ASC PYD filaments (Lu et al., 2014, 2015; Morrone et al., 2015; Shen et al., 2019). The cryo-EM structure reveals the molecular assembly of the Pyrin domain in the filament using asymmetric type I, II and III interfaces, similar to other filament-forming PYDs, despite low sequence conservation of only 22% identity. Particularly electrostatic interactions in the NLRP3^PYD^ as well as polar and hydrophobic contacts facilitate the assembly of individual PYDs into the right-handed helical filament. Importantly, we show at the protein level that the NLRP3^PYD^ filament can act as a nucleation seed for ASC filament elongation. This transition is exclusively mediated on one end of the NLRP3^PYD^ filament, which we defined as the ‘B-end’ according to the interface sites Ib, IIb and IIIb that assemble to one surface and form this filament end. The directionality of the homotypic PYD filament transition determines the growth direction of the ASC speck and sets the molecular basis for understanding these micrometer-large signaling assemblies.

The contribution of individual residues of the NLRP3^PYD^ to form filaments was confirmed by site-directed mutagenesis with recombinant proteins and in cells. The charge reversal mutations introduced at the interface sites each abrogated the ordered formation of a filament (Figure 3). Yet, some NLRP^PYD^ mutants such as E15R or M27E still showed a strong tendency for aggregation, indicating their ability for homomeric interactions, albeit in an uncoordinated manner. The two CAPS mutants D19H and H51R appear of particular interest, both showing a tendency for an increased filament polymerization activity compared to wild type NLRP3^PYD^. This is seen not only in negative-stain EM but also by their ability to nucleate ASC in cells. It suggests that a well-coordinated PYD/PYD transition between the sensor and adaptor molecule is critical for optimal inflammasome function. Our studies provide a biochemical basis of NLRP3 hyperactivity in patients carrying gain-of-function mutations in the PYD domain of NLRP3. Aspartate-19 is part of interface III but not involved in any salt bridge or hydrogen bond formation (Figure 1). The side chain points towards the filament’s hollow cylinder, most closely facing Glu46 of an adjacent molecule. One might speculate that the mutation to histidine enhances the interaction to the carboxylic groups of Glu46, forming a new hydrogen bond in this interface. Histidine-51 instead contributes only a small van-der-Waals interaction to interface Ia with a portion of 28 out of 500 Å^2^ buried surface area. Its exchange to a larger arginine could form a new interaction network by reaching out with a hydrogen bond to Gln45 of the interface IIIa from a neighboring subunit and a salt-bridge to Glu15 of the same molecule. Interestingly, AIM2, which has indeed an arginine at this position, contains instead a charge repulsive lysine at the position of Gln45 and a smaller aspartate at the position of Glu15, prohibiting such interaction network. Our observation that the ability of the NLRP3^PYD^ to form filaments could even be increased by the CAPS mutations D19H and H51R underlines how tightly balanced the interplay between autoinhibition and effector function is in inflammasomal regulation. Thus, the NLRP3^PYD^ does not seem to be evolutionarily designed for the firmest filament formation, but could represent a compromise for multiple molecular states, including the homotypic transition to another molecule as ASC^PYD^.

Regulation of inflammasome formation also occurs at the level of post translational modifications (Yang et al., 2017). Phosphorylation of NLRP3 at S5 was shown to prevent NLRP3^PYD^ oligomerization in HEK293T cells (Stutz et al., 2017). The NLRP3^PYD^ filament determined here shows that the hydroxyl group of S5 participates in the intra-strand interface type I through hydrogen bonding with D31. Introducing a bulky phosphorylation at S5 containing a negative charge repulsive to D31 is likely to disrupt the tight salt-bridge and hydrogen bonding interplay of this interface (Figure S6A). Conversely, acetylation of NLRP3 at residues K23 and K24 was shown to trigger aging-associated chronic inflammation through promoting NLRP3 inflammasome formation (He et al., 2020). Residues K23 and K24 participate in type I interface interactions in both, the NLRP3^PYD^ filament and the modelled NLRP3^PYD^ to ASC transition. Acetylation of NLRP3 residues K23 and K24 disrupts the salt-bridges formed with ASC residues D51 and E13 respectively, however it establishes a new network of hydrogen-bonds that might facilitate increased ASC to NLRP^PYD^ filament binding (Figure S6B).

The formation of filamentous structures in the cytosol is a very rare and typically tightly controlled process in mammalian cells. The by far best-characterized examples are the cytoskeleton structures of actin and microtubules. These two structures are reversible and highly regulated by numerous factors that control nucleation, elongation, capping, severing and crosslinking (Goodson and Jonasson, 2018; Pollard and Borisy, 2003). Moreover, both are nucleotide-binding proteins and their polymerization activity depends on nucleotide and magnesium concentrations (O’Brien et al., 1990; Swenson et al., 2014). The direction of filament growth is indeed essential for understanding the dynamics of the filament structure and its dependency on (ligand-)concentration and steady-state equilibria. In contrast to the dynamic cytoskeleton, filament formation of the NLRP3^PYD^ and ASC is irreversible. Both filaments form readily with recombinant protein in a concentration and pH-dependent manner, but once filaments are formed, they do not disassemble upon dilution. A close interplay of on and off rates at the A and B ends, similar to the barbed and pointed end of the actin filament, leading to a treadmilling mechanism, does not seem to exist in the PYD assembly of NLRP3 and ASC, as far as we know today. Instead, the formation of the NLRP3^PYD^ and ASC filaments appear as a ‘point of no return’ that will ultimately lead to cell death and pyroptosis.

It is remarkable that ASC only assembles at one site of the NLRP3^PYD^ filament, which defines the growth direction of the ASC speck. From the EM data presented here, we cannot differentiate if there is a structural constraint, *e.g.*, in form of a steric clash, that prohibits the assembly of the ASC^PYD^ to the A-end of the NLRP3^PYD^ filament, or if the on-rate of the a-side PYD interfaces of monomeric ASC for binding to the B-end of the NLRP3^PYD^ filament is so much in favor compared to binding in the opposite direction that virtually only one form of the homotypic domain transition occurs. It is indeed tempting to speculate that only the interaction kinetics determines the homotypic domain transition directionality and that also the filament growth direction, *e.g.*, of the NLRP3^PYD^ alone, is unidirectional and determined by the binding kinetics. Finally, it can also be envisioned, that topological constraints allow only one growth and PYD transition direction. The NLRC4 protein was shown to form a disc-like structure in an assembly of up to eleven subunits in its active state (Diebolder et al., 2015; Hu et al., 2015; Tenthorey et al., 2017; Zhang et al., 2015). The large LRR domain mainly contributes to the planar assembly of the disc, while the death-domain coalesce in the central hub. Also, for NLRP3 a disc-like assembly with participation of the serine/threonine kinase NEK7 was predicted previously (Sharif et al., 2019). Moreover, the NLRP3 basic region 131–147 following the NLRP3^PYD^ (3-94) was shown to associate with lipid membranes (Chen and Chen, 2018). While it is not clear if active NLRP3 launches the inflammasome formation from a membrane surface, it is well conceivable that filament growth is only possible in one direction if this disc-like structure associates on a membrane surface.

Knowing the direction of the homotypic NLRP3 to ASC PYD transition and piecing together recent findings from other laboratories, we generated a molecular model of an ASC speck (Figure 7). A comprehensive analysis of ASC filament formation, serving as a signal amplification mechanism for inflammasomes, was performed previously (Dick et al., 2016). It was shown that assembly of the ASC speck involves the oligomerization of ASC^PYD^ into filaments while cross-linking these filaments is mediated by the CARD (Dick et al., 2016; Schmidt et al., 2016). Accordingly, ASC mutants with a non-functional CARD only assemble filaments but do not form specks. While NLRP3 is the sensor and its PYD the nucleation seed for inflammasome formation, ASC is the adaptor that transmits as a filament elongation factor the information to the effector protein Caspase-1 (Lu and Wu, 2015; Tapia-Abellán et al., 2021). This cascade ultimately leads to ASC specking, Caspase-1 activation, IL-1β maturation and GSDMD cleavage. We generated a model of eleven NLRP3–NEK7 complexes assembling into the nucleation seed for ASC filament elongation. The ASC^PYD^ forms the stem of the elongating ASC filament while the CARDs assemble as a bud onto the surface of the stem. In the model, we placed 32 ASC molecules with the N-terminal PYD arranged in the filament structure, as determined by Lu et al. (Lu et al., 2014). The C-terminal CARD again is assembled in a new filament structure (Li et al., 2018) or in any of the 20 conformers determined by NMR spectroscopy (de Alba, 2009). The length of the 23 residue-linker between the PYD and CARD of ASC easily allows for the formation of two rings of the four-leafed CARD filament on the ASC^PYD^ stem (Figure 7A). This ‘bud of CARDs’ tethered to the PYD stem acts as a new nucleation seed for Caspase-1 recruitment and subsequent Casp^CARD^ filament formation (Lu and Wu, 2015). Of note, the ASC^CARD^ filament diameter is ∼7 nm (Li et al., 2018) and thus 1.5 nm smaller than the ASC^PYD^ filament. The ASC^CARD^ filament structure assembles through a left-handed one-start helical symmetry with about 3.6 subunits per turn (Li et al., 2018). The following transition of the ASC^CARD^ to the Caspase-1^CARD^ marks the last step in this domino-type NLRP3^PYD^:ASC^PYD^-ASC^CARD^:Casp^CARD^ queue. Remarkably, there are at least 3.5-fold more Caspase-1 molecules found in the ASC speck than ASC molecules (Lu et al., 2014), giving rise to the assembly of 120 Caspase-1 proteins in our model. Onto the three ASC^CARD^ buds with four to ten subunits each, 54 to 66 Caspase-1 molecules were modelled, either as full-length protein or in its cleaved form with CARD, p20, and p10 subunits (Boucher et al., 2018) (Figure 7B). Interestingly, the transition zone from the active NLRP3^PYD^ filament seed to the ASC^PYD^ filament in a functional inflammasome can be as short as one ASC^PYD^ filament ring as shown for the ASC (E80R) mutant that retained the ability to induce cell death (Dick et al., 2016). These data nicely confirm our experimentally determined directionality of NLRP3^PYD^ to ASC filament growth, as the ASC E80R mutation is located on the b-side of the PYD interface surfaces, allowing for the transition of the NLRP3^PYD^ filament B-end to the ASC^PYD^ A-end, whereas subsequent ASC filament elongation is prohibited by this mutation. Accordingly, the ASC^CARD^ bud for a functional inflammasome can be as short as only six ASC subunits. The model of an ASC speck shown here displays the minimal assembly of NLRP3, ASC, and Caspase-1 proteins in a rough 1:3:10 stoichiometry. While this model holds for an approximate 80 nm length, the ASC^CARD^ mediated cross-linking (Dick et al., 2016; Schmidt et al., 2016) of multiple such assemblies may generate the micrometer-like structure seen in fluorescence microscopy for the ASC speck (Fernandes-Alnemri et al., 2007; Glück et al., 2021; Kagan et al., 2014).

**Figure 7.**
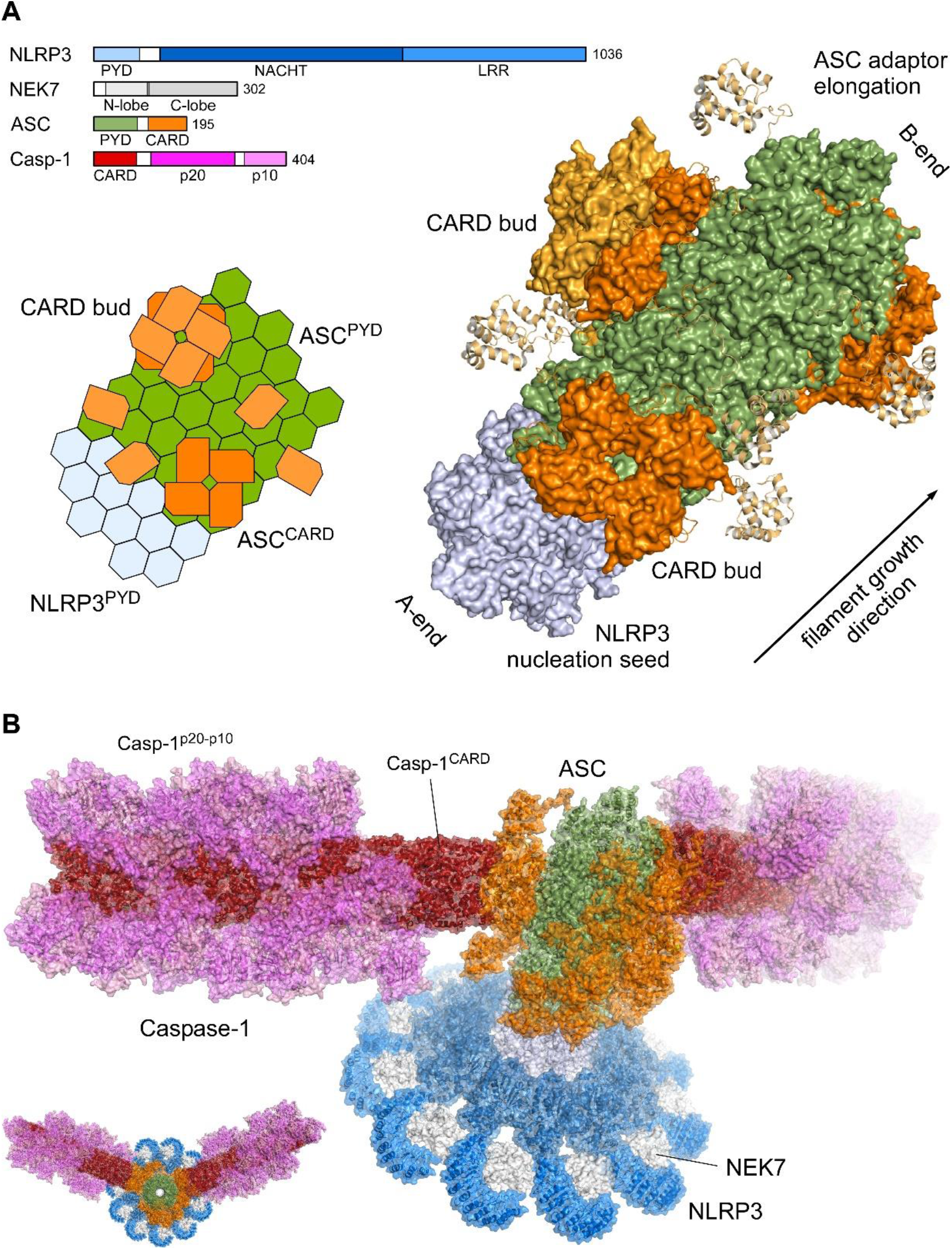
Molecular model of an ASC speck. (A) Cartoon and atomistic model of the NLRP3^PYD^–ASC filament elongation. The molecular domain architecture of human NLRP3, NEK7, ASC and Caspase-1 is displayed in bar diagrams. The NLRP3^PYD^ nucleation seed (light blue) and the elongation to the ASC^PYD^ filament (green) are shown with the ASC^CARD^ (orange) assemble on the stem. The linker between the PYD and CARD domains in ASC allow small ASC^CARD^ nucleation seed (CARD buds) of one up to three filament rings. (B) Molecular model of an ASC speck. The NLRP3–NEK7 complex forms the disc-like assembly (blue, grey) of the active NLR with the PYD filament in the center core (light blue). The ASC^PYD^ stem (green) with the CARD buds (orange) constitute the adaptor to Caspase-1 effector filament with the Casp-1^CARD^ filament (red) and the zymogen p20-p10 dimer (violet and pink) attached. The structures used to generate this model assembly were: NLRP3–NEK7 complex (6NPY(Sharif et al., 2019)), NLRP3^PYD^ filament (this structure), ASC^PYD^ filament (3J63 (Lu et al., 2014)) with an overlay with the ASC full length NMR structure on the PYD subunit (2KN6 (de Alba, 2009); 20 models), ASC^CARD^ buds (assembly of 1 or 2 ASC^CARD^ rings on the ASC^PYD^ stem) (6N1H (Li et al., 2018)), ASC^CARD^–Casp1^CARD^ transition (7KEU (Robert Hollingsworth et al., 2021)), Casp1^CARD^ filament (5FNA (Lu et al., 2016)), and Casp1-zymogen p20-p10 dimer (3E4C (Elliott et al., 2009)).

The NLRP3^PYD^ filament structure and the determination of the transition directionality to ASC filament elongation reveal the molecular basis for NLRP3-mediated ASC specking. Our observations have implications for the possible interference with antibodies or small molecules at the filament growing site. Capping the inflammasome at the filament growth direction could be an effective mechanism to block these higher-order signalosomes and lead to new therapeutic intervention in inflammation.

## METHODS

### NLRP3^PYD^ cloning, expression, and purification

Human NLRP3 (UniProt accession number Q96P20, residues 3-110, C108S) was cloned into a pGEX-4T1 expression vector providing an N-terminal glutathione S-transferase (GST) tag followed by a tobacco etch virus (TEV) protease cleavage site. The C108S mutation was introduced in order to prevent disulfide bond formation with residue C8, which was observed in one of two chains in the crystal structure (Bae and Park, 2011) of NLRP3^PYD^. The construct was transformed and expressed in *E. coli* cells, strain BL21(DE3), by growing the culture at 37°C to an OD_600_ of 0.8 and inducing with 0.3 mM isopropyl β-D-1-thiogalactopyranoside (IPTG) o/n at 20°C. Cells were collected by centrifugation and lysed by sonication in a lysis buffer containing 20 mM HEPES (pH 7.5), 150 mM NaCl and 0.5 mM TCEP. The cell lysate was centrifuged at 20,000 x g for 30 min, and the supernatant was administered to a pre-equilibrated GSTrap column using an ÄKTA prime FPLC system (GE Healthcare). The column was washed with 10 column volumes of lysis buffer, and the protein was eluted in the same buffer supplemented with 15 mM L-Glutathione. The affinity-purified protein was TEV-cleaved o/n at 4°C and subsequently subjected to gel filtration on an equilibrated Superdex 75 gel filtration column 16/600 (GE Healthcare). Monomer fractions were pooled and concentrated to 1 mg/ml, corresponding to a concentration of 79 µM, and filament formation was induced with fresh protein by incubation at 37°C overnight.

Protein variants carrying mutations in the PYD-PYD assembly interfaces (R7E, E15R, K23E/K24E, M27E, R43E, D60R, R80E, R81E) and CAPS mutants (D19H, D31V, H51R, A77V) were generated in the NLRP3^PYD^ (3-110, C108S) construct using the mega primer mutagenesis method. The NLRP3^PYD^ interface and CAPS mutant proteins were expressed and purified as GST-fusion proteins following the same procedure as described above. For kinetic measurements, the generation of monomeric, soluble NLRP3^PYD^ included an additional dialysis step after TEV-protease cleavage into a buffer containing 50 mM glycine (pH 3.8) and 150 mM NaCl. The dialyzed protein was further purified on a pre-equilibrated Superdex 75 gel filtration column (GE Healthcare), and fractions containing monomeric soluble NLRP3^PYD^ were concentrated to 1 mg/ml, snap-frozen in liquid nitrogen, and stored at −80°C.

### Preparation of recombinant ASC

Full-length human ASC, followed by a TEV protease cleavage site and mCherry, was cloned with *Nde*I/*Xho*I sites at the 5’ and 3’ ends, respectively, into a pET-23a expression vector providing a C-terminal hexa-histidine tag. This construct was transformed and expressed in *E. coli* cells, strain BL21(DE3), by growing the culture at 37°C to an OD_600_ of 0.8 and induced with 0.1 mM IPTG for 4 h at 37°C. Cells were collected by centrifugation and lysed by sonication in lysis buffer A containing 20 mM Tris (pH 8.0), 500 mM NaCl and 5 mM imidazole. Cell lysates were centrifuged at 20,000 x g for 30 min, and the pellet was dissolved in buffer A supplemented with 4 M Gdn-HCl for 1 h at 4°C. Subsequently, the suspension was centrifuged at 20,000 x g for 30 min, and the supernatant was administered onto a pre-equilibrated HisTrap column using an ÄKTA Prime FPLC system (GE Healthcare). The column was washed with 10 column volumes of solubilization buffer, supplemented with 20 mM imidazole, and the protein was eluted in the same buffer supplemented with 300 mM imidazole. The pooled elution fractions’ pH was decreased to 3.8 and dialyzed against 50 mM glycine buffer (pH 3.8) and 150 mM NaCl. The protein was further purified on a pre-equilibrated Superdex 75 gel filtration column (GE Healthcare). Fractions containing monomeric, soluble ASC-mCherry protein were pooled, concentrated to 1 mg/ml, aliquoted, snap-frozen in liquid nitrogen, and stored at −80°C.

### Negative-stain EM

For negative-stain EM, 4 µl of protein sample was applied onto a glow-discharged copper grid coated with a continuous carbon layer (PLANO). The protein sample was incubated for 1 min before blotting away the excess sample with a filter paper. The sample coated side of the grid was then washed by dipping it into three individual 20 µl drops of protein buffer with a blotting step in between. Finally, the sample was negatively stained with 2% uranylformiate for 30 s, the excess sample was blotted away, and the grid was airdried. Negative-stained EM girds were imaged using a JEOL JEM-2200FS transmission electron microscope (TEM) operating at 200 kV.

### Cryo-EM sample preparation, data collection of NLRP3^PYD^ filaments and NLRP3^PYD^ to ASC-mCherry filament transitions

To enhance filament binding, R1.2/1.3 Cu 300 grids (Quantifoil) were glow discharged and coated with graphene oxide (0.2 mg/ml in ddH_2_O), followed by coating with poly-L-Lysin (1 mg/ml in ddH_2_O) prior to sample application. 4 µl of polymerized NLRP3^PYD^ (1 mg/ml) or NLRP3^PYD^:ASC-mCherry transitions (molar ratio 10:1) supplied with 0.01 % of the surfactant octyl maltoside, were then applied onto the freshly pretreated grids. Samples were blotted for 3-4 s at 80% humidity and 20°C and subsequently plunge-frozen in liquid ethane using an EM GP blotter (Leica Microsystems). Grids containing NLRP3^PYD^ filaments were imaged using a Krios Titan TEM (ThermoFisher), operated at 300 kV and equipped with a modified Falcon 2 direct electron detector enabling frame acquisition. Grids containing the NLRP3^PYD^:ASC-mCherry transition filaments were imaged with a Krios Titan TEM equipped with a Falcon 3 direct electron detector. For NLRP3^PYD^, 3000 frame movies with a total dose of 60 e^-^/A and 70 frames each were collected at a defocus range of −1 to −2.5 µm. For NLRP3^PYD^:ASC-mCherry, 1668 frame movies with a total dose of 60 e^-^/A and 40 frames each were collected in counting mode at a defocus range of −1.8 to −3 µm.

### Cryo-EM data processing and model building of the NLRP3^PYD^ filament

Cryo-EM data processing was done in RELION3 (He and Scheres, 2017). Frame movies were aligned using RELION’s own implementation of the MOTIONCORR-2 algorithm. Motion-corrected but non-dose-weighted micrographs were used to determine CTF parameters using CTFFIND 4.1. 724 high-quality micrographs were chosen for further processing, and the start and end coordinates of filaments were manually identified using the EMAN2 interface (Tang et al., 2007). In total, 100,821 particles were extracted in RELION from the motion-corrected and dose-weighted micrographs with a box size of 220 pixels assuming a helical rise of 14 Å. Particles were subjected to reference-free 2D classification, which yielded 13 2D classes showing high-resolution features accounting for a total of 60,333 particles. This subset was selected for further processing. An initial model was calculated based on the ASC^PYD^ filament structure (Lu et al., 2014) (PDB: 3J63). For this, the atomic model was first converted into a simulated electron density, low-pass filtered to 8 Å, and then symmetrized assuming a helical twist of 54° and an axial rise of 14 Å. 3D refinement was then performed using the same values as the initial search parameters for the helical symmetry. 3D alignment parameters were used to re-extract centered particle images from the micrographs. These were subjected to an intermediate round of 2D classification, which yielded ten classes with high-resolution features accounting for 26,172 particles.

Next, a subsequent round of 3D refinement was performed using these particles. For this, the map generated from the 60,333 particles was used to calculate a solvent mask. Helical parameters converged to a helical twist of 54.44° and an axial rise of 14.16 Å and yielded a resolution of 3.7 Å. Next, CTF refinement and Bayesian polishing were performed. A final round of 3D refinement produced a map at a resolution of 3.6 Å with a helical twist of 54.88° and an axial rise of 14.32 Å. Chain A of the dimeric NLRP3^PYD^ crystal structure (Bae and Park, 2011) (PDB: 3QF2) served as initial model for model building of the NLRP3^PYD^ filament in coot(Emsley et al., 2010). Real space refinement was done in Phenix (Adams et al., 2010). The final model encompasses residues 3-94 with clear densities of most side chains seen in the electron density map (Figure S1F). A multiple sequence alignment of all fourteen human NLRP PYDs displays the degree of sequence conservation in correlation to the secondary structure (Figure S7).

### Determination of NLRP3^PYD^ filament ASC-mCherry elongation directionality by cryo-EM

To determine the directionality of NLRP3^PYD^ filament seeded elongation by ASC-mCherry, we recorded a cryo-EM dataset of NLRP3^PYD^:ASC-mCherry transitions. The 1668 recorded movies were aligned using RELION’s own implementation of the MOTIONCORR-2 algorithm. Motion-corrected, non-dose-weighted micrographs were subjected to CTF-estimation using CTFFIND 4.1. In total, 423 micrographs that allowed a clear distinction of individual transition events were chosen for further processing. Start and end coordinates were always picked in the same direction relative to the filament transition site. The start coordinate was located at the transition distant, and the end coordinate was located at the transition proximal end of the generated particle box. In total 19,351 particles were extracted in RELION with a box size of 220 pixels and used for reference-free 2D classification. Four 2D classes (29, 44, 47, 48) showed high-resolution features and a characteristic saw-tooth pattern. Only these “good classes” were used in the following procedure. The directionality of the class averages with respect to the PYD filament structure was determined by manual superposition of the electron density onto the class averages. The directionality of the class average was then flagged by overlaying arrows onto the class average.

To identify directionality of the NLRP3^PYD^–ASC^PYD^ transition, we extracted the x- and y-coordinates, psi- and psi_prior_ angles from each of the particles that constituted the four good classes. The directionality-flagged class averages were then aligned onto the extracted x,y positions in the CTF- and motion-corrected micrographs and rotated according to the extracted psi-angle (Figure S4). The process was repeated for 100 micrographs, and in each case, the B-end of the class average pointed toward the ASC-mCherry filament transition. Also, we noted that since the particle picking was always performed towards the ASC-mCherry transition, the psi and psi-prior angles of the particles were always very similar. Figure S4 shows histograms of the Δpsi angles (psi_prior_–psi) of all particles in the good classes. Due to our described picking procedure of picking towards the ASC-mCherry transition, the histograms are centered around 0°. This means that RELION did not change the overall direction of any of the particles during the formation of the class averages. Any filaments with a reversed directionality would have led to peaks at 180° in the histogram. Very few instances of +360° or −360° occurred for filaments that were almost horizontal in the micrograph and thereby aligned along the x-axis (*e.g.*, 179.2°-(−178.7°) = 357.9°), which is however in agreement with the identified transition directionality.

### NLRP3^PYD^ filament polymerization monitored by DLS

Prior to the kinetic analysis of NLRP3^PYD^ polymerization, a sample containing monomeric, soluble NLRP3^PYD^ (50 mM glycine pH 3.8, 150 mM NaCl, 0.5 mM TCEP) was rapidly thawed, centrifuged at 13,000 rpm for 5 min and filtered through a 0.1 µm syringe filter (Whatman). The protein concentration was adjusted to 0.6 mg/ml using sample buffer and filament formation was induced by adjusting the sample to pH 8.0 through addition of 3 M Tris (pH 8.0) to a final concentration of 60 mM. Filament polymerization was monitored by batch dynamic light scattering (DLS) with a DynaPro Nanostar (Wyatt) instrument. Data were acquired at 25°C in a time course experiment of 100 min in 60 s intervals by averaging three runs of 20 s until a plateau of filament polymerization was reached.

For polymerization analysis of single-point mutants, recombinant NLRP3^PYD^ and mutant proteins were purified in a buffer containing 20 mM HEPES (pH 7.5), 150 mM NaCl and 0.5 mM TCEP as described above, and SEC fractions containing monomeric recombinant protein were pooled and concentrated to 0.6 mg/ml. Filamentation was induced by incubation at 25°C for 1.5 h, and the polymerization status of the samples was subsequently monitored by batch DLS. Data were acquired by averaging three runs of 20 s. Three technical replicates of each experiment were performed.

### Polymerization of NLRP3^PYD^ and point mutants analyzed by negative-stain EM

In addition to DLS the polymerization behavior of wild type NLRP3^PYD^ and point mutants was analyzed by negative-stain EM. Here, GST-NLRP3^PYD^ or GST-NLRP3^PYD^ point mutants at a concentration of 1 mg/ml were subjected to TEV-protease cleavage (molar ratio TEV-protease: GST-NLRP3^PYD^ of 1:25) for 1.5 h at 25°C and subsequently analyzed by negative-stain EM as described above.

### ASC specking experiments

5000 HeLa cells stably over-expressing ASC-mTurquoise were seeded and transfected in duplicates with increasing amounts (0, 3.1, 6.3, 12.5, 25, 50, 100 or 200 ng) of plasmids encoding wild type and point mutant NLRP3 (1-95)-mCitrine fusion proteins. 36 hours after transfection, cells were fixed, and nuclei were stained using a PBS solution containing both paraformaldehyde (4%) and DRAQ5 (1:2000). Ten images per well were taken using filter sets to detect CFP (*i.e.*, filter set 47 from Zeiss: excitation BP 436/20, beamsplitter FT 455, emission BP 480/40) and DRAQ5 (*i.e.*, filter set 50 from Zeiss: excitation BP 640/30, beamsplitter FT 660, emission BP 690/50) using the 20x objective of a Zeiss observer Z1 microscope. Images were analyzed using CellProfiler 2.2.0 software to count nuclei and ASC specks. For each well, the ratio ASC specks/nuclei was calculated. Ratios extracted from 20 independent images (10 images per well, conditions in duplicates) were averaged to calculate the final ratio ASC speck/nuclei for every condition.

### NLRP3^PYD^ seeding ASC filament elongation assays

To induce filament formation for homotypic PYD transition experiments, 50 µl of monomeric NLRP3^PYD^ protein (50 mM glycine pH 3.8, 150 mM NaCl, 0.5 mM TCEP) at a concentration of 50 µM were adjusted to pH 8.0 by adding 1 µl of 3 M Tris (pH 8.0) to a final concentration of 60 mM. The solution was incubated for 3 min at 25°C, allowing for the formation of short NLRP3^PYD^ filaments. Monomeric soluble ASC-mCherry (50 mM glycine pH 3.8, 150 mM NaCl, 0.5 mM TCEP) was added to the NLRP3^PYD^ sample to reach a molar ratio of 1:100 (ASC-mCherry to NLRP3^PYD^), and the sample was incubated for 5 min. This step was repeated two more times at increasing ASC volumes to reach consecutive molar ratios of ASC-mCherry to NLRP3^PYD^ of 1:50 and 1:25, respectively. After each addition and subsequent incubation with the ASC-mCherry protein, the protein solution was applied onto EM grids, negatively stained, and imaged using a JEOL JEM-2200FS microscopy to visualize binary complex assemblies containing ASC-mCherry filaments topping on NLRP3^PYD^ filament seeds.

For the cryo-EM procedure the titration protocol to generate NLRP3^PYD^:ASC-mCherry filament transitions was adjusted to receive longer initial NLRP3^PYD^ filaments that could be used for subsequent particle extraction. The boxed particles were used for determining the directionality of ASC-mCherry elongation on NLRP3^PYD^ filaments. Here, 50 µl of monomeric NLRP3^PYD^ protein (50 mM glycine pH 3.8, 150 mM NaCl, 0.5 mM TCEP) at a concentration of 50 µM were adjusted to pH 8.0 as described above, and the solution was incubated for 6 min at 25°C. Monomeric soluble ASC-mCherry (50 mM glycine pH 3.8, 150 mM NaCl, 0.5 mM TCEP) was added to the NLRP^PYD^ sample to reach a molar ratio of 1:50 (ASC-mCherry to NLRP3^PYD^) followed by 5 min incubation. In two consecutive steps, the molar ratio of ASC-mCherry to NLRP3^PYD^ was increased to 1:25 and 1:10, respectively, each followed by 5 min incubation. The sample of the last incubation step was used for cryo-EM sample preparation.

### Molecular modeling of an ASC speck

A model of an 11-mer NLPR3–NEK7 complex was created by separate structural alignment of the NACHT and LRR-NEK7 subunits of (PDB-ID 6NPY) (Sharif et al., 2019) to each subunit of the disc-like NLRC4 structure (3JBL) (Zhang et al., 2015). This ring-shaped structure was placed at the A-end of a 12 subunit NLRP3^PYD^ filament (our structure, 7PZD). At its B-end, this filament was elongated by 32 subunits of an ASC^PYD^ filament (3J63) (Lu et al., 2014) and the transition between both filaments was energy minimized by GROMACS (Van Der Spoel et al., 2005). The NMR ensemble of full-length ASC (2KN6) (de Alba, 2009) was split into 20 individual structures. Each ASC^PYD^ in the filament was overlayed with a randomly selected NMR structure to create a “full-length” model (with respect to the ASC polypeptide) of the ASC filament. Due to the 24 amino acid-linker between the ASC^PYD^ and ASC^CARD^ domains, it was possible to model eight subunit-large ‘buds’ of an ASC^CARD^ filament at three different positions of the ASC^PYD^ filament. The arrangement of the subunits in these buds was modelled by using the ASC^CARD^ filament structure (6N1H) (Li et al., 2018) as a template. A superposition of the ASC^CARD^ to CASP1^CARD^ transition structure (7KEU) (Robert Hollingsworth et al., 2021) onto the buds created the transition to the CASP1^CARD^ filament (5FNA) (Lu et al., 2016). Finally, structural models of the CASP1-zymogen (p20-p10 dimer; 3E4C) (Elliott et al., 2009) were placed at each junction of two CASP1^CARD^ molecules in the filament.

### Data availability

The cryo-EM structures have been deposited in the Electron Microscopy Data Bank (EMDB) with accession numbers EMD-13727 for the NLRP3^PYD^ filament. The atomic coordinates and structure factors have been deposited in the protein databank (PDB) with accession number 7PZD. All other data that support the findings of this study are available from the corresponding author upon request.

## Supplemental Information

Supplemental information can be found online at https://

## Acknowledgments

We thank the research center caesar, Bonn, and the Rudolf-Virchow-Zentrum at Universität Würzburg for EM measurement time. M.G. and E.L. are funded by the Deutsche Forschungsgemeinschaft under Germany’s Excellence Strategy–EXC2151–390873048. This work was supported by a grant from the Else Kröner-Fresenius-Stiftung to M.G. (2014_A203).

## Author contributions

I.V.H. purified proteins and performed biochemical experiments. I.V.H. and H.B. prepared cyro-EM samples. I.V.H. and E.B. processed cryo-EM data. I.V.H. and G.H. built the model. I.V.H., G.H. and M.G. interpreted the data. I.V.H. and A.K. established the NLRP^PYD^ seeding assay for ASC filament elongation. J.F.R.A. performed ASC specking cell assays that E.L. supervised. I.V.H. and M.G. wrote the paper. All authors contributed to editing the manuscript and support the conclusions.

## Supplemental figures

**Figure S1.**
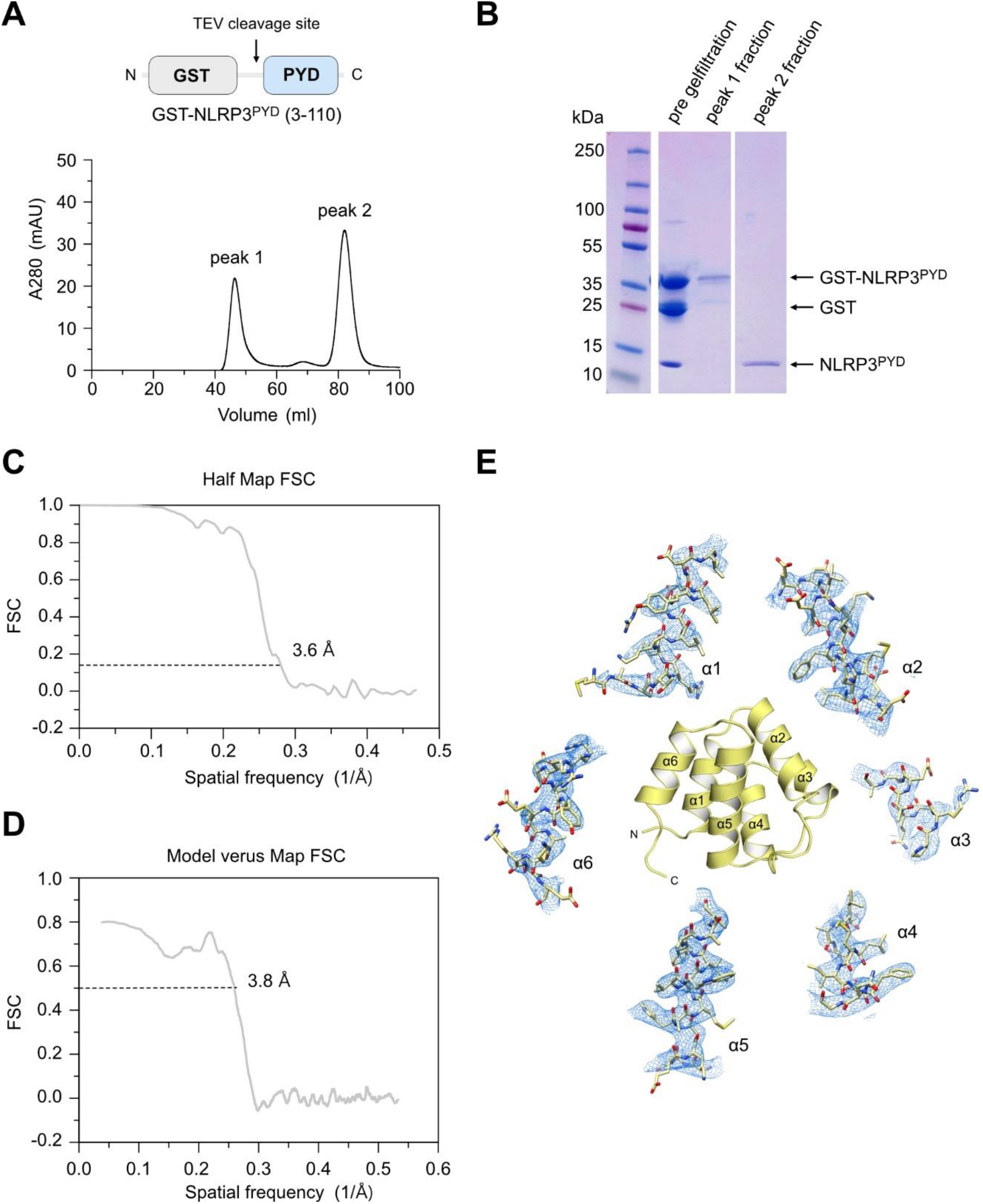
Cryo-EM structure determination of the NLRP3^PYD^ filament, related to Figure 1. (A) Construct design and gel filtration of human NLRP3^PYD^ protein. Recombinant NLRP3^PYD^ is expressed as GST fusion protein with a TEV cleavage site before the PYD. After expression in *E. coli* and glutathione-affinity purification, some protein elutes near the void fraction (peak 1) whereas the larger fraction remains monomeric (peak 2). (B) SDS PAGE analysis of peak 1 and peak 2 fractions, indicating purification to homogeneity for NLRP3^PYD^ in peak 2. (C) Gold standard Fourier shell correlation (FSC) of the final model. (D) Resolution measurement of the NLRP3^PYD^ filament. FSC of model versus map. (E) Fitting of NLRP3 helices α1-α6 into the final electron density map, showing clear densities of most side chains.

**Figure S2.**
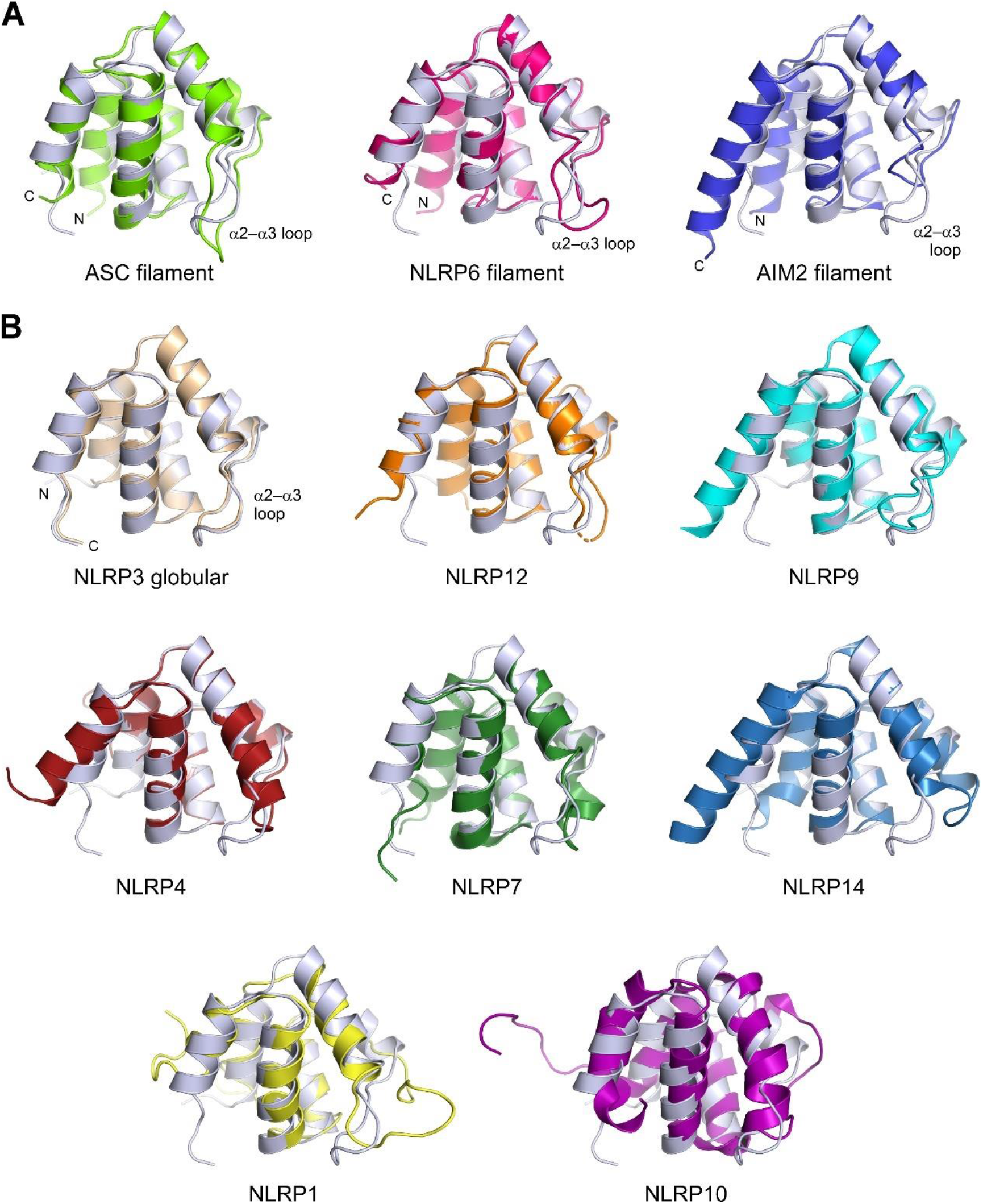
Comparison of the NLRP3^PYD^ filament structure with filament forming PYDs and other globular PYDs, related to Figure 1. (A) Overlay of a single PYD subunit from the NLRP3^PYD^ filament structure (light blue) with other filament forming PYDs ASC (3J63) (Lu et al., 2014): rmsd value 1.370 Å over 51 atoms; NLRP6 (6NCV) (Shen et al., 2019): 1.022 Å over 62 atoms; and AIM2 (6MB2) (Lu et al., 2015): 1.376 Å over 68 atoms. Largest deviations are seen within the α2–α3 loop that is two residues shorter in NLRP6 and six residues shorter in AIM2 compared to NLRP3 and ASC. (B) Overlay of a single PYD subunit from the NLRP3^PYD^ filament structure with monomeric PYDs NLRP3 (3QF2, chain A) (Bae and Park, 2011): 0.537 Å over 84 atoms; NLRP12 (5H7N) (Jin et al., 2017): 0.535 Å over 71 atoms; NLRP9 (6Z2G) (Marleaux et al., 2020): 0.778 Å over 68 atoms; NLRP4 (4EWI) (Eibl et al., 2012): 0.661 Å over 65 atoms; NLRP7 (2KM6) (Pinheiro et al., 2010): 1.133 Å over 69 atoms; NLRP14 (4N1L) (Eibl et al., 2014): 0.762 Å over 62 atoms; NLRP1 (1PN5) (Hiller et al., 2003): 1.096 Å over 61 atoms; and NLRP10 (2M5V) (Su et al., 2013): 4.486 Å over 73 atoms.

**Figure S3.**
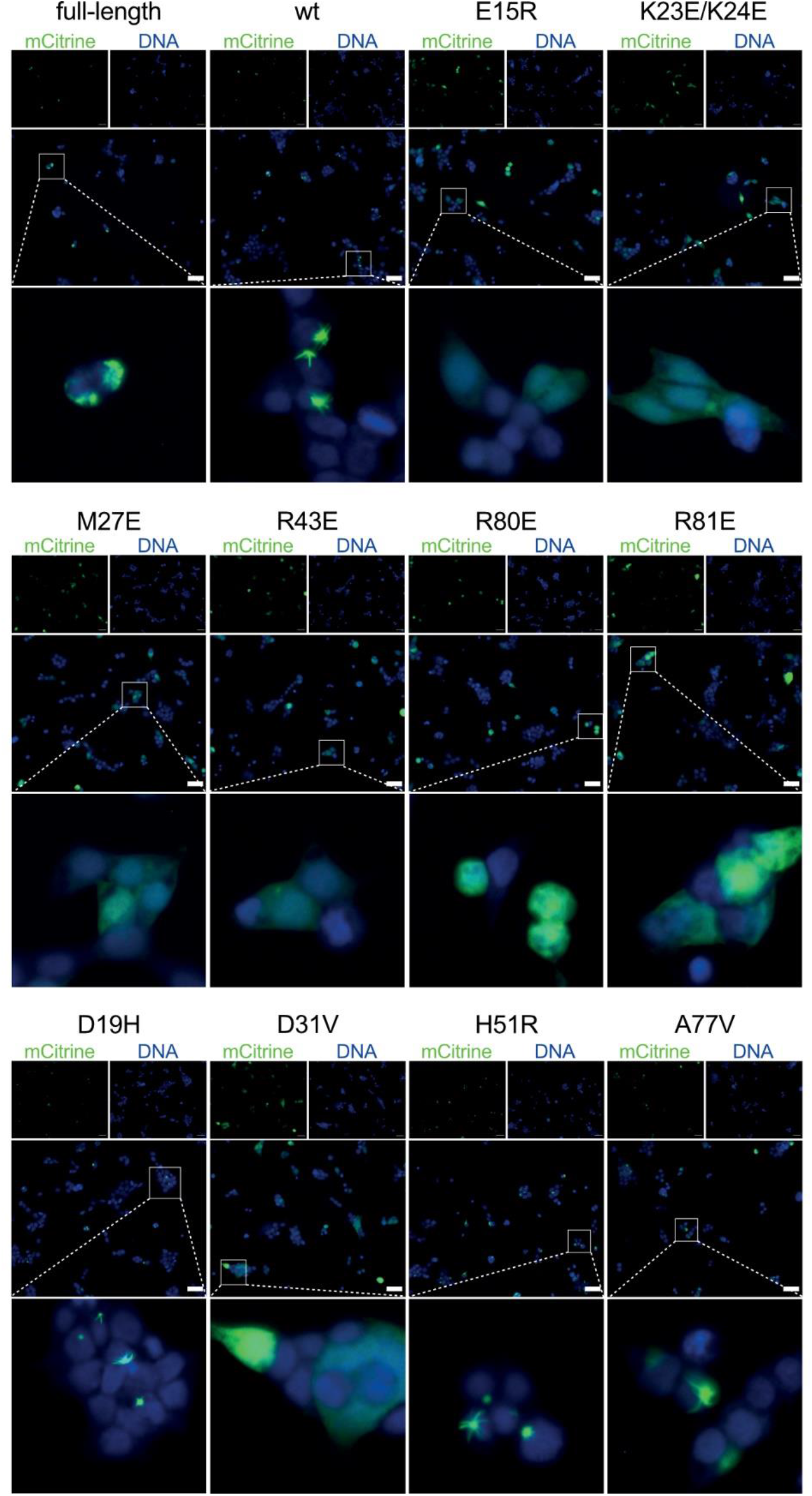
ASC speck formation of wild-type and mutant NLRP3 in cells, related to Figure 3. Images of HeLa cells stably expressing ASC-mTurquoise transfected with full-length wild type NLRP3-mCitrine, wild type NLRP3^PYD^-mCitrine, and ten different mutants. Scale bar, 50 µm.

**Figure S4.**
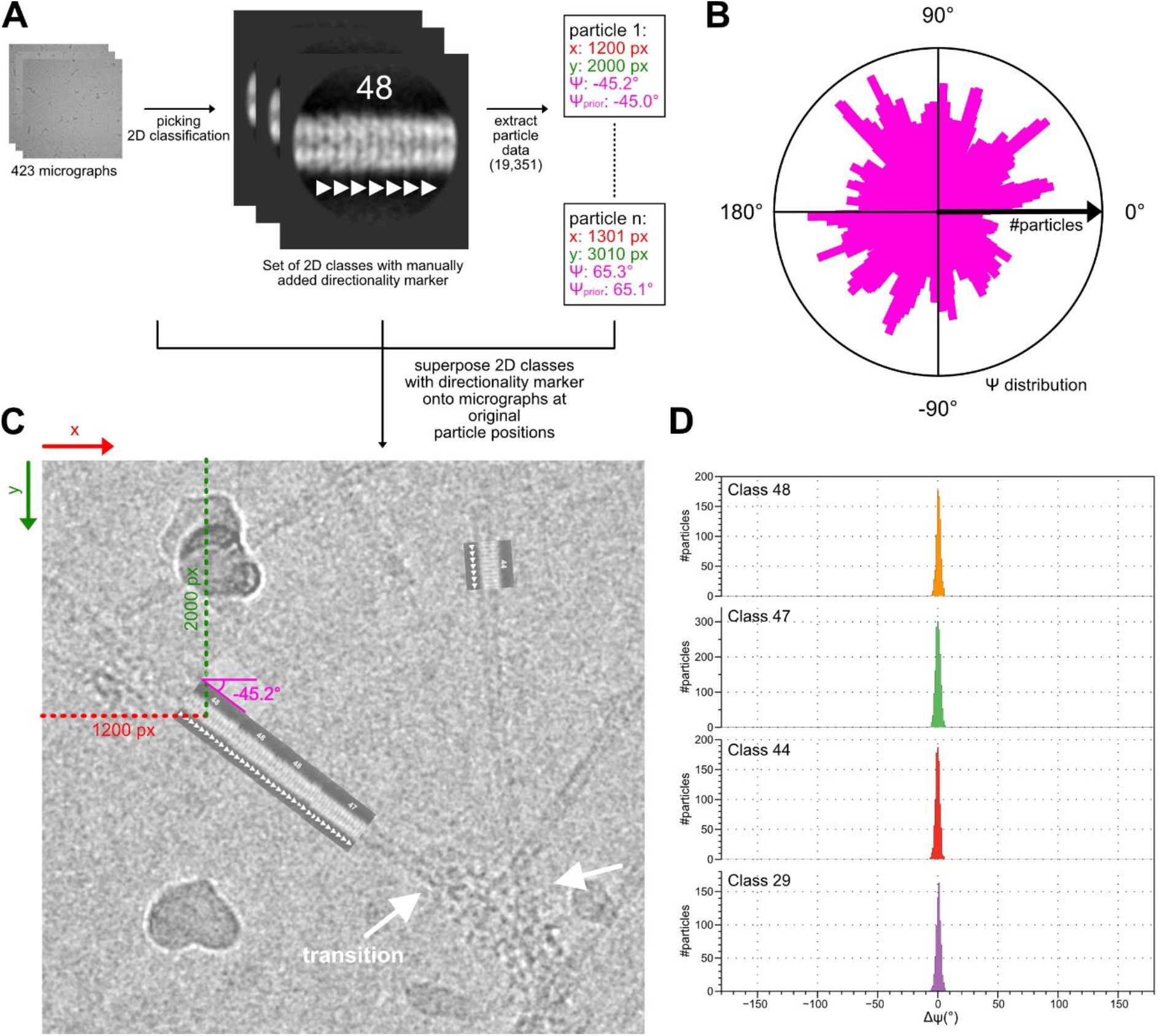
Determination of the directionality of NLRP3^PYD^ to ASC^PYD^ transitions, related to Figure 5. (A) NLRP3^PYD^ filament particles with visible transitions to ASC filament elongation were manually picked with RELION (He and Scheres, 2017) and grouped into 2D classes. The best 2D classes that showed a clear sawtooth pattern (*e.g.*, class 48) were selected and a directionality indicator (white triangle) was manually added to the class averages. The alignment parameters (x, y, psi_prior_ and psi) of the underlying particles were extracted. (B) A polar histogram showing the distribution of psi angles (*i.e.*, the direction of the filaments) in the dataset. (C) To determine the directionality of the filaments in the micrographs, the individual class averages were aligned onto the filaments according to the previously extracted alignment parameters (see A). In every case, the manually added directionality indicators ended up pointing in the same direction, towards the transition between the two filaments (white arrows). (D) Histogram, showing the distribution of the psi_prior_–psi angles for all particles in the different classes. The distribution is centered around zero because the manual picking was always done in the direction of the transition.

**Figure S5.**
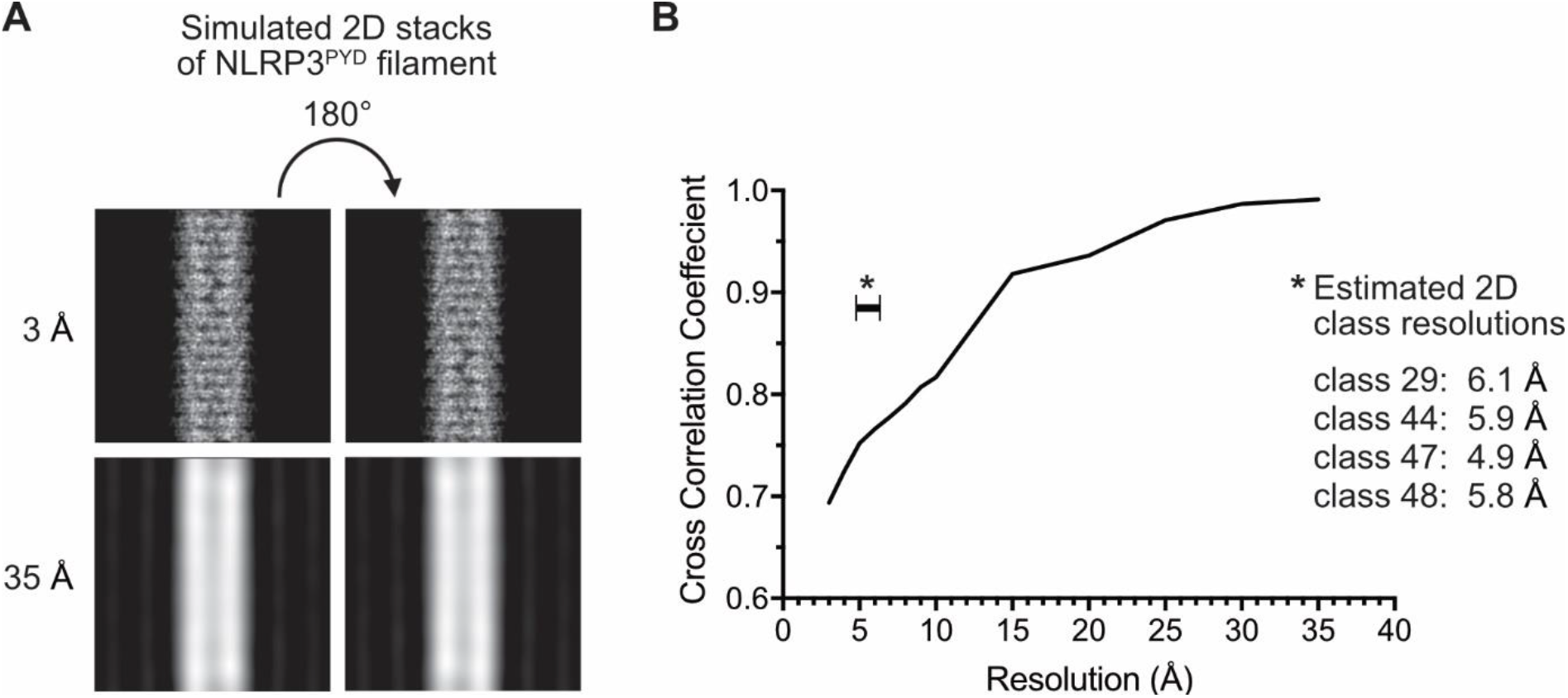
Discrimination of NLRP3^PYD^ filament directionality at different resolutions, related to Figure 5. (A) The NLRP3^PYD^ filament structure was simulated at different resolutions ranging from 3 to 35 Å. Simulated maps were 2D projected and the cross-correlation coefficients calculated between the original 2D projection and the same projection rotated by 180°. (B) Cross-correlation coefficients of the simulated 2D projections of the NLRP3^PYD^ filament were plotted as a function of resolution. Resolutions of the 2D classes (e.g., classes 29, 44, 47 and 48) used for the determination of NLRP3^PYD^ to ASC transition directionality are indicated (*).

**Figure S6.**
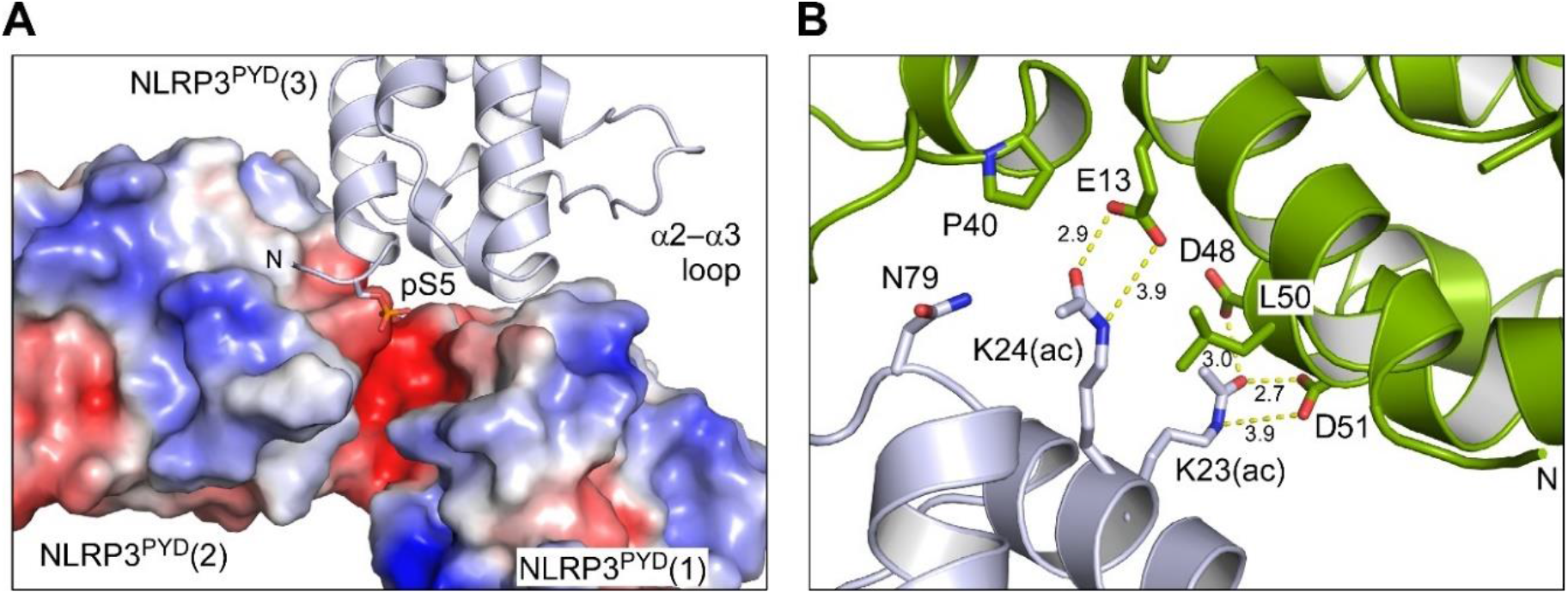
Compatibility of NLPR3^PYD^ post translational modifications with filament formation, related to Figure 6. (A) Molecular modelling reveals that phosphorylation of Ser5 will lead to a steric clash with Asp31 of another NLRP3^PYD^ molecule, and an electrostatic repulsion with a negatively charged surface patch. The electrostatic surface potential is displayed from −4 *k*_B_*T* (red) to +4 *k*_B_*T* (blue). (B) Modelling the acetylation of lysines 23 and 24 in NLRP3^PYD^, shown here in the transition interface to ASC^PYD^, reveals that this post translational modification is well compatible with the formation of a hydrogen bond network that involves the carboxylic groups of Glu13, Asp48, and Asp51 of the opposing PYD subunit. These three residues of the a-side of interface I are identical in the NLRP3^PYD^.

**Figure S7.**
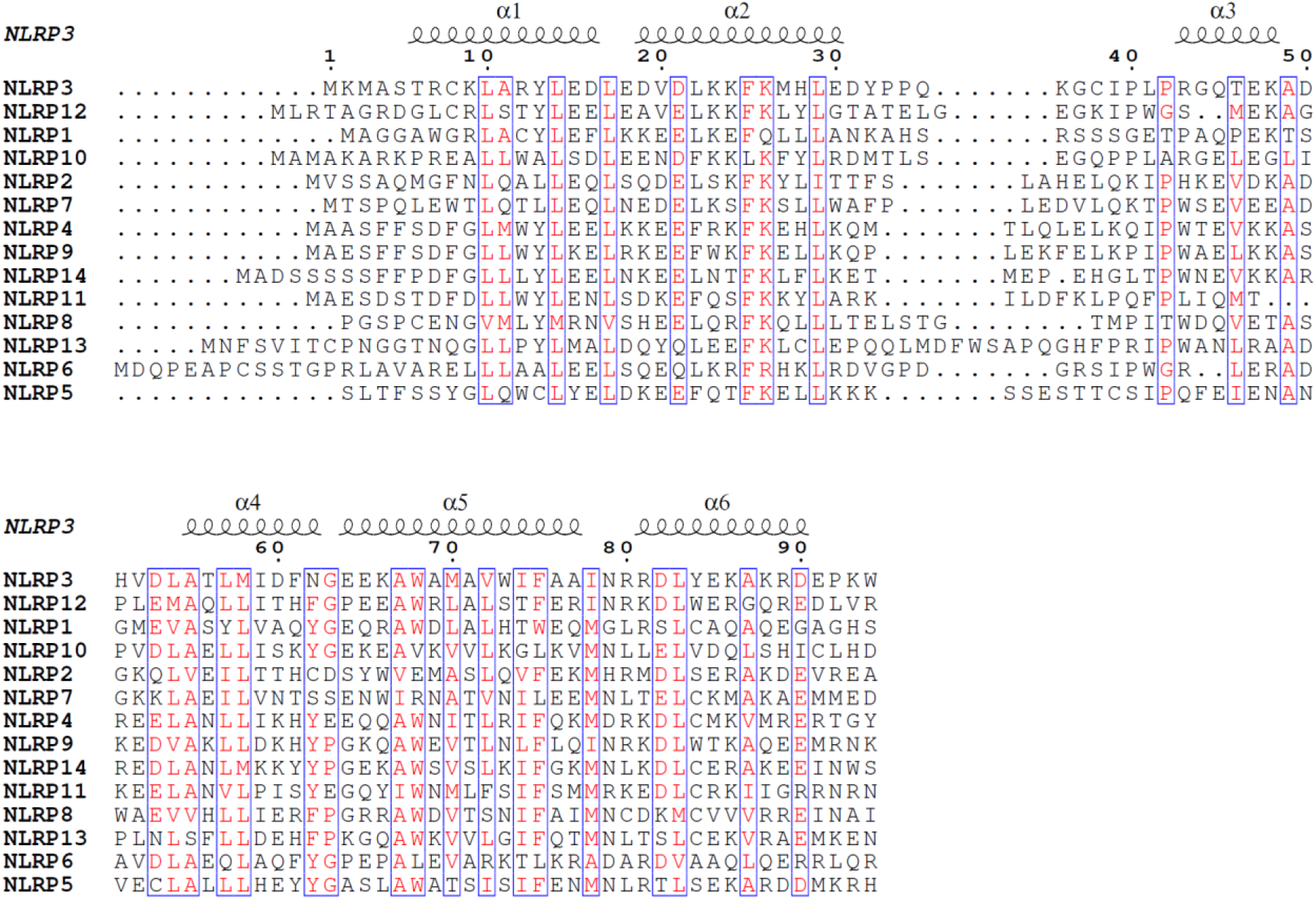
Sequence alignment of the fourteen human NLRP Pyrin domains, related to Figure 1. Display of all fourteen human NLRP PYDs arranged according to the sequence conservation to NLRP3. The secondary structure elements of the NLRP3^PYD^ in the filament structure determined here are shown on the top. The multiple sequence alignment was performed with Clustal Omega and the sequence conservation is colored and boxed using ESPript (Robert and Gouet, 2014).

**Table S1.** Cryo-EM data collection, refinement and validation statistics, related to Figures 1 and 5.

|  | **NLRP3<sup>PYD</sup>** | **NLRP3<sup>PYD</sup>:ASC** |
| --- | --- | --- |
| **Data collection and processing** |  |  |
| Magnification | 59,000 | 47,000 |
| Voltage (kV) | 300 | 300 |
| Electron exposure (e<sup>-</sup>/Å<sup>2</sup>) | 60 | 60 |
| Defocus range (μm) | -1 to -2.5 | -1.8 to -3 μm |
| Pixel size (Å) | 1.067 | 1.77 |
| Symmetry imposed | helical | helical |
| Initial particle images (no.) | 100,821 | 19,351 |
| Final particle images (no.) | 26,172 |  |
| Map resolution (Å) at FSC threshold 0.143 | 3.6 |  |
| **Refinement** |  |  |
| Initial model used (PDB code) | 3QF2 |  |
| Model resolution (Å) at FSC threshold 0.5 | 3.8 |  |
| Map sharpening *B* factor (Å<sup>2</sup>) | -117.282 |  |
| **Model composition** | for 18 chains |  |
| Non-hydrogen atoms | 13,644 |  |
| Protein residues | 1,656 |  |
| **R.m.s. deviations** |  |  |
| Bond lengths (Å) | 0.010 |  |
| Bond angles (°) | 1.306 |  |
| **Validation** |  |  |
| MolProbity score | 1.58 |  |
| Clashscore | 3.31 |  |
| Poor rotamers (%) | 1.27 |  |
| **Ramachandran plot** |  |  |
| Favored (%) | 94.44 |  |
| Allowed (%) | 5.56 |  |
| Disallowed (%) | 0 |  |
| **Data accessibility** |  |  |
| EMDB accession code | EMD-13727 |  |
| PDB accession code | 7PZD |  |

